# Multi-modal pooled Perturb-CITE-Seq screens in patient models define novel mechanisms of cancer immune evasion

**DOI:** 10.1101/2020.09.01.267211

**Authors:** Chris J. Frangieh, Johannes C. Melms, Pratiksha I. Thakore, Kathryn R. Geiger-Schuller, Patricia Ho, Adrienne M. Luoma, Brian Cleary, Shruti Malu, Michael S. Cuoco, Maryann Zhao, Meri Rogava, Lila Hovey, Asaf Rotem, Chantale Bernatchez, Kai W. Wucherpfennig, Bruce E. Johnson, Orit Rozenblatt-Rosen, Dirk Schadendorf, Aviv Regev, Benjamin Izar

## Abstract

Resistance to immune checkpoint inhibitors (ICI) that activate T cell mediated anti-tumor immunity is a key challenge in cancer therapy, yet the underlying mechanisms remain poorly understood. To further elucidate those, we developed a new approach, Perturb-CITE-seq, for pooled CRISPR perturbation screens with multi-modal RNA and protein single-cell profiling readout and applied it to screen patient-derived autologous melanoma and tumor infiltrating lymphocyte (TIL) co-cultures. We profiled RNA and 20 surface proteins in over 218,000 cells under ~750 perturbations, chosen by their membership in an immune evasion program that is associated with immunotherapy resistance in patients. Our screen recovered clinically-relevant resistance mechanisms concordantly reflected in RNA, protein and perturbation effects on susceptibility to T cell mediated killing. These were organized in eight co-functional modules whose perturbation distinctly affect four co-regulated programs associated with immune evasion. Among these were defects in the IFNγ-JAK/STAT pathway and in antigen presentation, and several novel mechanisms, including loss or downregulation of *CD58*, a surface protein without known mouse homolog. Leveraging the rich profiles in our screen, we found that loss of *CD58* did not compromise MHC protein expression and that *CD58* was not transcriptionally induced by the IFNγ pathway, allowing us to distinguish it as a novel mechanism of immune resistance. We further show that loss of *CD58* on cancer cells conferred immune evasion across multiple T cell and Natural Killer cell patient co-culture models. Notably, CD58 is downregulated in tumors with resistance to immunotherapy in melanoma patients. Our work identifies novel mechanisms at the nexus of immune evasion and drug resistance and provides a general framework for deciphering complex mechanisms by large-scale perturbation screens with multi-modal singlecell profiles, including in systems consisting of multiple cell types.

## INTRODUCTION

Molecular circuits form the basic driving force from genotype to phenotype across all levels, from cells to tissues to entire organisms. Circuits in cells process diverse signals, make appropriate decisions, and orchestrate physiological responses to these signals. Diseases arise from circuit malfunctions: one or more components are missing or defective; a key module is over- or under-active. One key approach to chart circuits and understanding their function is pooled perturbation screens. Because pooled screens require a cell-autonomous readout, they were historically limited to either simple phenotypes (*e.g.*, viability) or to specific markers. However, this required choosing markers *a priori*, and cannot further distinguish between hits, thus giving limited information about the ways in which different genes impact the same ultimate phenotype through different pathways (*1*).

The recent development of methods like Perturb-seq (*2–5*) combined pooled genetic perturbation screens, where through single cell RNA-seq (scRNA-seq) the perturbation is read as an RNA barcode along with the full RNA profile of the cell, as a rich readout. This provides a powerful platform to link specific perturbations with cell phenotypes and enables identification of converging and unique cell programs impacted by different perturbations, as well as allows screening in systems where cells are in different cell states (*2*). However, many relevant phenotypes are functionally best understood at protein level, and are reflected differently at the RNA and protein level, thus requiring a modification to prior Perturb-Seq schemes.

A key case in point is using screens to enhance our understanding of resistance to immune checkpoint inhibitors (ICIs). Antibodies targeting CTLA-4 or the PD-1/PD-L1 axis release tumor-mediated immune inhibition and enable T-cell mediated killing of tumor cells (*6*). While ICIs produce durable responses in some patients across diverse cancer types, most patients either do not benefit or develop resistance over time. Understanding mechanisms of resistance is therefore a key challenge and opportunity. Genomic profiling of DNA and RNA directly from patient tumors have begun to shed light on some of the genetic and epigenetic mechanisms underlying intrinsic and acquired resistance to ICI. Genetically, these include mutations in beta-2-microglobulin (*B2M*) that cause loss of MHC Class I presentation and downregulation of the antigen presentation machinery, and mutations in the IFNy-JAK/STAT pathway, which result in impaired response to T cell mediated anti-tumor activity (*7–9*). At the level of cells states, using single-cell RNA-seq (scRNA-seq) of patient tumors, we previously identified a signature present in a subset of malignant cells in metastatic melanoma tumors predictive of intrinsic resistance to ICI (*9*), and can be reversed by CDK4/6 inhibitors and re-sensitize tumors to combination therapy with ICI in patient cell co-cultures and mouse models (*9*). However, genetic signals only account for a minority of cases, and signatures like the ICR signature consist of dozens of genes, many of which may play specific roles, and would require systematic investigation. In parallel, perturbation screens in in mice or engineered human cell lines have identified both known mutations of immune evasion from patients involving the IFNy-JAK/STAT pathway, and putative mechanisms involving chromatin regulators (*10*), TNFα signaling (*17*), *SOX4* (*12*), *PTPN2* (*13*), and *APLNR* (*10*). However, the clinical significance of these observations remains unclear. Moreover, previous screens focused on malignant cell viability (by enrichment or depletion of the perturbations) and could not distinguish mechanism-of-action, as would be reflected by molecular phenotypes in the RNA or protein level. Moreover, the models used, based on either animal models or established cell lines, may be less faithful to human tumors. Recent advances in cellular models should allow us to study co-cultures of malignant cells derived directly from patient tumors along with their *ex vivo* expanded autologous tumor infiltrating lymphocytes (TILs) (*14*). Such models would better reflect human tumor biology, but have not yet been used in a screening context.

Here, we develop Perturb-CITE-seq, an extension of Perturb-Seq that combines scRNA-seq profiling and epitopes sequencing (CITE-seq) (*15*) of single-cell surface proteins under perturbations (*16*), and apply it to dissect critical interactions at the tumor-immune synapse. We perform pooled Perturb-CITE-Seq screens of the ICR program genes in a unique patient-derived malignant-TIL co-culture model, targeting 248 genes of the ICR signature (744 targeting guides) and profiling single-cell transcriptomes and 20 surface proteins in >218,000 cells. We inferred a computational model relating eight co-functional modules to four co-regulated gene programs, which recapitulated the landscape of clinically relevant resistance mutations in the IFNγ-JAK/STAT pathway and antigen-presentation machinery, related them to the genes and biological processes they regulate, and identified novel driver pathways of immune evasion in perturbation, RNA and protein space. In particular, downregulation or loss of *CD58* in malignant cells conferred resistance to T cell mediated killing of melanoma cells by TILs, which our model showed reflects a new mechanism of immune resistance, that does not act by impacting the MHC Class I or II complexes, is not a component of the IFNγ-JAK/STAT module, and is not induced by IFNγ signaling. As there is no mouse homolog for *CD58*, such mechanisms can only be identified in human systems (*10*). Using multiple patient-derived co-culture models we show that loss of *CD58* on cancer cells confers resistance to both T cell and Natural Killer (NK) cell mediated killing. Finally, *CD58* is strongly downregulated in melanoma patients with resistance to immunotherapy, supporting its role in immune evasion and drug resistance. Our work presents new rich functional predictions of mechanisms of immunotherapy resistance, and validation of a novel resistance mechanism that is orthogonal to previously described mechanisms in patients and provides a broadly applicable experimental and analytical framework for Perturb-CITE-Seq.

## RESULTS

### A patient-derived co-culture model for viability and Perturb-CITE-Seq screens of resistance to T cell mediated killing

To enable a systematic and functional evaluation of the contribution of genes in the ICR signature to resistance to T cell mediated killing, we designed two types of CRISPR/Cas9 loss of function (KO) screens using a human tumor-immune co-culture model (*9*) under increasing immune pressure (**Figure 1A**): a viability screen, to determine the impact of perturbation on T cell-mediated killing (**Figure 1H**) and a Perturb-CITE-seq screen, to decipher the underlying circuitry (**Figures 1I-1J**).

**Figure 1.**
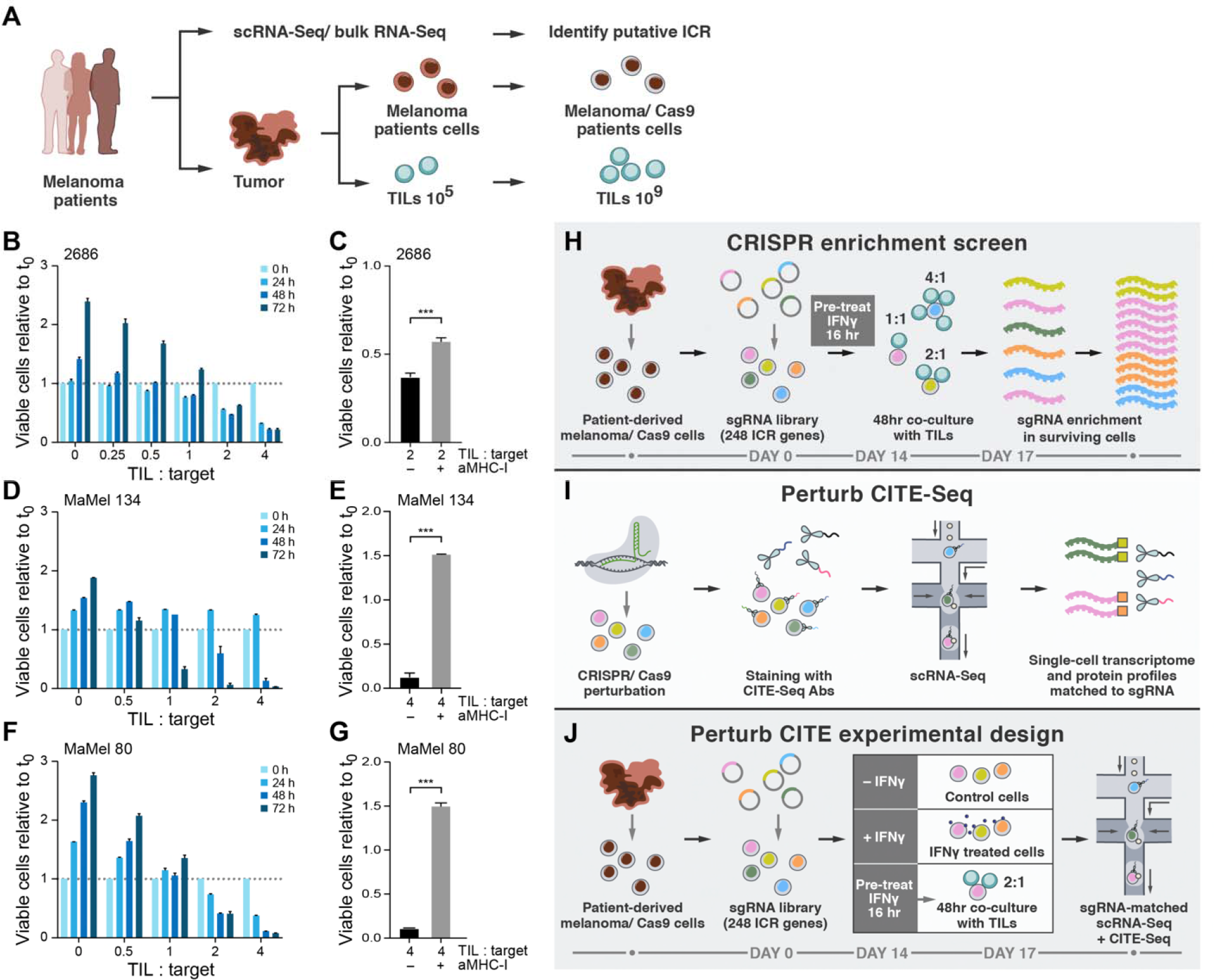
Perturb-CITE-Seq to study tumor intrinsic mechanisms of T cell resistance using patient derived co-culture models. **(A)** Patient derived cell models and programs. Resected melanomas (left) were profiled by bulk and scRNA-seq (top) to identify an immune resistance program (ICR) (*9*). Melanoma (brown, middle) and tumor infiltrating lymphocytes (TILs, blue, bottom) cells were grown from patients’ tumors; melanoma cells were transduced to express Cas9 and TILs were expanded to yield sufficient numbers for screening (Methods). (B)-(G) Optimization and validation of co-culture system for TIL mediated killing of melanoma cells. (B)(D)(F) Time and dose-dependent killing of melanoma target cells by autologous TILs in three patient derived co-culture models. Ratio of viable cancer cells (y axis, relative to t_0_) at different TIL : cancer cell ratios (x axis) at different time points (color legend) after pre-treatment of target cells with 1 ng/ml IFNγ for 16 hours using TILs without prior restimulation in each patient-derived co-culture model (panels). Experiment was performed in triplicates in each of two independent experiments. (C)(E)(G) Target cell killing depends on MHC-I. Ratio of viable cancer cells (y axis, relative to t_0_) after 48 hours in a 4:1 TIL and cancer cell co-culture with cancer cells pre-treated with 1ng/ml IFNγ for 16 hours and TILs without prior restimulation, in the absence or presence of MHC-I blocking antibodies (x axis). *, p<0.05; **, p<0.005; ***, p<0.001, *t*-test. Experiment was performed in triplicates in each of two independent experiments. (H) Viability screen design. (I) Perturb-CITE-seq approach. (J) Perturb-CITE-seq screens to characterize regulators of melanoma immune evasion.

We first established autologous co-culture models of melanoma cell lines and *ex vivo* expanded TILs from multiple patients (**Figure 1A; Methods**). The expanded TILs were exclusively CD8+ T cells (**Figures S1B and S1D)** or a mixed population of CD8+ and CD4+ T cells (**Figure S1C)**. Using a miniaturized optical platform (**Figure S1A; Methods**) and improved methods of T cell activation to reduce bystander killing (**Figures S1B-S1H; Methods**), co-cultures of T cells and cancer cells resulted in dose- (TIL:cancer cell ratio) and time-dependent cancer cell killing (**Figures 1B, 1D, and 1F**). Tumor cell lysis was highly reproducible, specific to T cell receptor (TCR)/MHC Class I interactions unique to autologous pairs, and absent in allogenic controls (**Figure S1H**), and rescued with MHC blocking antibodies (**Figures 1C, 1E, and 1G**).

Next, for both viability and Perturb-CITE-Seq pooled screens, we established a pooled library of 744 single-guide (sg)RNA targeting 248 genes with putative roles in immunotherapy resistance (ICR library, **Supplementary Data Table 1**), cloned into a modified CROP-seq (*4*) vector (**Methods**). For the viability screen, we included 758 control guides (379 non-targeting and 379 targeting intergenic regions), and for the Perturb-CITE-seq screen we had 74 control guides (37 non-targeting and 37 intergenic). We transduced patient-derived melanoma cells that stably expressed high-activity Cas9 (**Figure S2A**) with the ICR library at MOI 0.1 to achieve a high rate of single-guide transductions, with ~1,000 cells/guide (**Figures S2B-S2D**), cultured the cells for 14 days, and then either co-cultured them with a range of TIL doses (1:1, 2:1 or 4:1 ratios), treated them with IFNγ (no co-culture), or maintained them in culture media alone (control). For the viability screen (**Figure 1H**) survivor cells of each condition were collected after 48 hours of co-culture (day 17 after library transduction) for gDNA isolation, sequencing and identification of enriched perturbations (**Methods**). (We also sequenced the gDNA from the control population on days 7 and day 14 to identify perturbations targeting “essential” genes that are underrepresented prior to exposure to TILs.) For the Perturb-CITE-Seq screen, we collected surviving cells from each of the three conditions (with co-cultures performed at 2:1 ratio) and performed Perturb-CITE-seq (**Figure 1J**) using droplet based scRNA-seq, with each perturbation represented by ~100 cells (*2*).

### A co-culture CRISPR-Cas9 viability screen identifies known and novel drivers of immune evasion in human tumor cells

The strongly enriched perturbations in the robust (**Figures S3A-S3C**) viability screen spanned the entire known landscape of clinically established mechanisms of resistance to immunotherapies in patients (*7–9*) and is consistent with previous murine screens (*10–13*). First, to recognize “essential” genes independent of T cell mediated killing, we identified sgRNAs that are depleted by day 7 and day 14 prior to any treatment or co-culture (**Figures 2A and S3B-S3D; Supplementary Data Table 2**). These included expected essential genes such as *MYC, ATP1A1, EIF2S3, RPSA*, and *TUBB.* On day 14, we treated cells with low-dose IFNγ or control media for 16 hours, followed by either media only or co-culture with TILs for 48 hours at 1:1, 2:1 or 4:1 TIL:tumor cell ratios (**Figure 1H**). The screens were highly reproducible across triplicate samples (**Figures 2B, 2C, S3B, and S3C**), and co-culture showed dose-dependent lysis of cancer cells with 30.96%, 58.35 % and 75.0% killing at 1:1, 2:1 and 4:1 ratios, respectively (**Figure S3A**). The strongly enriched perturbations (**Methods**) that conferred resistance to T cell mediated killing included guides targeting *B2M, HLA-A, JAK1, JAK2, STAT1, IFNGR1* and *IFNGR2* (**Figures 2D, 2E**, **S3E, and 3F**; **Supplementary Data Table 2**), recovered all the known, clinically established mechanisms of resistance to immunotherapy in patients (*7–9*), and are consistent with previous screens in murine models (*10–13*).

**Figure 2.**
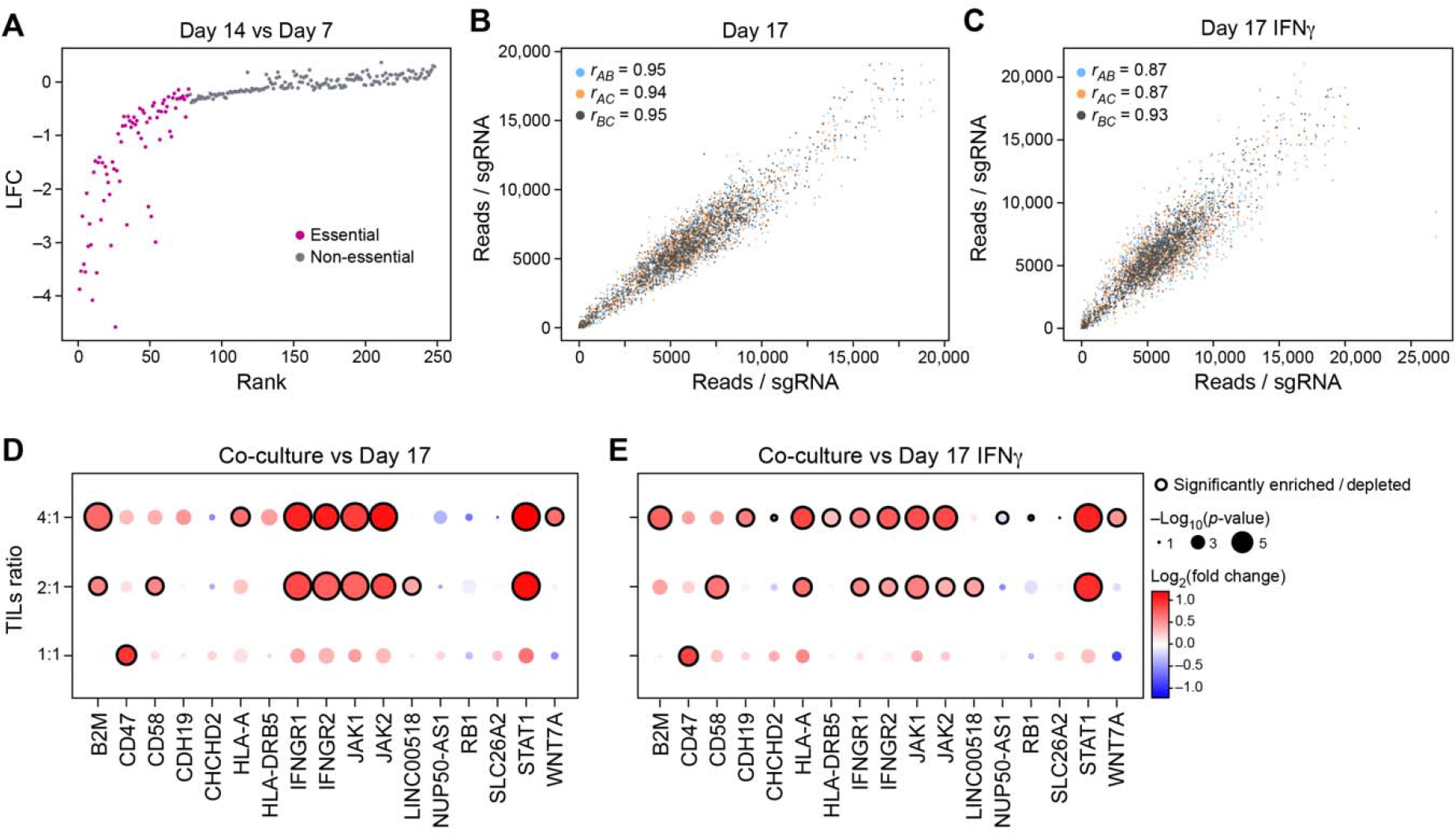
Identification of genes for evasion of TIL-mediated killing by CRISPR-Cas9 viability screen in patient-derived models exposed to increasing immune pressures. **(A)** Identification of essential gene unrelated to immune pressure. Change in abundance (negative log_2_ fold-change (LFC), y axis) of each sgDNA (dot) in day 14 *vs*. day 7 following lentivirus transduction, with guides ranked (*x* axis) by LFC value. Pink: called essential genes (**Methods**). **(B)(C)** High reproducibility of screen across triplicates. Number of reads detected (*x*, *y* axis) for each sgDNA (dots) when comparing each pair within triplicate experiments (color legend) in either control cells (B) and in IFNγ treated cells (C) on day 17. Pearson (*r*) correlation coefficients are noted in the color legend. (**D)(E**) Co-culture screen highlights role for IFN γ/Jak**-** STAT pathway and additional mechanisms. Log_2_(Fold Change) (dot color) and significance (-log_10_(*p-*value), dot dize (and significantly enriched/depleted cirled with black border), **Methods)** of genes (columns) whose sgDNA in tumor cells co-cultured with different doses of TILs (rows) was differentially enriched compared to control cells on Day 17 (**D**) or IFNγ-treated cells on day 17 (**E**).

The screen also identified additional genes that when targeted conferred resistance to T cell mediated killing, including *CD47, CD58, CDH19, WNT7A* and the long non-coding RNA *LINC00518.* Notably, all these perturbations were found to be enriched when compared to either IFNγ treated or to control cells, suggesting that these were specific to evading T cell mediated killing (**Figures 2D and 2E**). CD47 is a “don’t eat me” signal on tumor cells that interacts with SIRPa on phagocytic cells; and blockade of the CD47/SIRPa axis improves tumor control *in vivo* (*17*). Recent studies revealed that activated T cells also express SIRPa and that interaction with CD47 results in increased T cell activation and cytokine production (*1*), which may impact the potential clinical benefit of CD47-blockade (*18*). CD58 (also known as LFA3) is an understudied adhesion molecule, typically expressed on antigen-presenting cells, that binds CD2 on T cell and Natural Killer (NK) cells (*19*). The role of *CD58* in cancer is poorly understood. Notably, there is no known mouse homolog of *CD58*, emphasizing the value of performing such screens in human models (*10*). Overall, the viability screen recovered known immune evasion mechanisms in a dose-dependent manner, and identified several novel candidates. We next turned to our Perturb-CITE-Seq data to explore the organization of these in pathway along with the molecular processes they impact.

### Single cell RNA and protein profiles show coherent changes in RNA and protein levels for genes with functional impact in viability screen

Across all Perturb-CITE-Seq screens, we analyzed 218,331 high quality single-cell RNA and protein profiles across experimental conditions spanning the co-culture (73,114 cells), IFNγ (87,590), and control (57,627) conditions (**Figures 3A, 3E, and S4A-S4D; Methods**). We removed 805 contaminating CD8^+^CD28^-^ T cells using unsupervised clustering (**Figures S4A and S4B**; **Methods**). We first focused on the RNA and protein profiles in the context of the different conditions (irrespective of subsequent perturbations) to assess if their profiles can provide meaningful phenotypes in this system. Notably, embedding cells by either protein (**Figure 3A**) or RNA (**Figure 3E**) profiles separately differentiated treatment conditions in a similar way. Within each condition, both the cell cycle (G1/0, S, G2/M) (**Figure 3F**) and complexity (**Figure S4C**) impacted RNA profiles, revealing important covariates, which we address in later analyses. Notably, cell embedding by protein profiles did not similarly reflect the cell cycle or complexity (**Figures S4D and S4E**), and expression of some key immune proteins (*e.g.*, HLAs) was not affected by the cell cycle state (**Figures S4F-S4H**).

**Figure 3.**
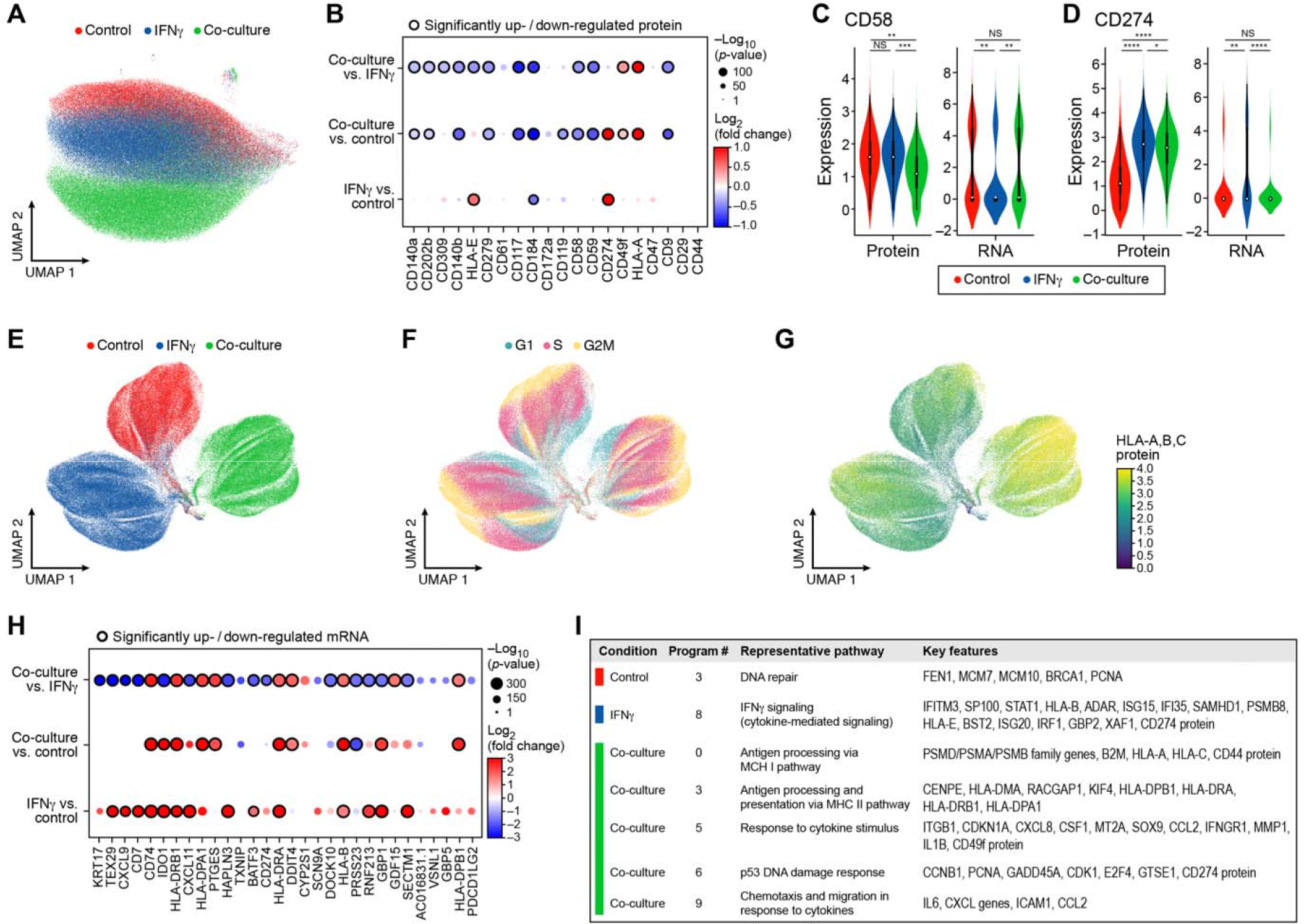
Single-cell protein and RNA profiles reveal regulation of genes and program involved in immune evasion. **(A)(B)** Distinct protein profiles across immune pressures highlight regulation of cell surface proteins whose genetic perturbation confers resistance to TIL mediated killing. (A) Uniform Manifold Approximation and Projection (UMAP) embedding of single cell CITE-antibody count profiles (dots) colored by condition (color legend). (B) Log_2_(Fold Change) (dot color) and significance (-log_10_(*p-*value)), dot size (and statistically significantly up/downregulated circled with black border), logistic regression model; Methods) between each pair of conditions (rows) of each cell surface protein (columns) measured by CITE-Seq. (C)(D) Regulation of CD58 and CD274 (PDL1) by culture conditions. Distribution of protein counts (y axis, left) or RNA (normalized expression, Methods) (y axis, right) for CD58 (C) and CD274 (D). **** P < 1^-10^, Welch’s t test. (E)-(G) Variation in RNA profiles across and within conditions captures cell cycle state and MHC-I protein expression. UMAP embedding of scRNA-seq profiles (dots) colored by condition (E), cell cycle phase signature (F), or HLA-A,B,C antibody expression level (G, color bar). (H)(I) RNA expression of key immune genes and programs is impacted by increased immune pressure. (H) Log_2_(Fold Change) (dot color) and significance (-log_10_(*p-*value)), dot size (and statistically significantly up/downregulated circled with black border) (*30*) between each pair of conditions (rows) of the RNA of select immune genes (columns) measured by scRNA-seq, and differentially expressed between conditions. (I) Gene programs identified by jackstraw PCA in each condition, representative enriched Gene Ontology processes, and select member genes.

Changes in both protein and RNA levels between conditions (quantified relative to matched isotype controls (*15*), **Supplementary Data Table 3, Methods**) reflected expected changes, including in genes that impacted T cell mediated killing in the viability screen. At the protein level, these included increase in HLA-A, HLA-E and PD-L1 (CD274) proteins in IFNγ-treated *vs.* control cells; a global increase in HLA-A protein in co-culture *vs*. control cells (**Figure 3B; Supplementary Data Table 4; Methods**), and, consistent with prior reports (*12*), induction of *CD49f*, an integrin associated with epithelial-to-mesenchymal transition (EMT), specifically in co-culture with TILs. Conversely, there was strong down-regulation of several proteins with potential roles in modifying the response to immunotherapies exclusively in co-culture conditions, including CXCR4 (CD184), c-KIT (CD117) and KDR (CD309) (*20, 21*) (**Figure 3B**). In particular, CD58, CD47, and IFNGR1 (CD119) protein levels were reduced in coculture, consistent with how their genetic KO in the viability screen increased survival of cancer cells in the context of T cell mediated killing (**Figures 3B**, **2D, and 2E**). Concordant with singlecell protein profiles, comparing RNA profiles between conditions also highlighted genes involved in antigen presentation, chemokines and immune modulators (**Figures 3G and 3H; Supplementary Data Table 4**). Only some of the protein-level differences between treatments were observed at the level of the corresponding transcript (**Figure 3C**), highlighting the importance of simultaneous surface protein characterization, as well as of global RNA profiling, rather than a single marker screen. Overall, our results suggest that genes that have a functional impact on susceptibility to T cell mediated killing (based on their genetic perturbation in the viability screen) are also regulated at the protein and RNA level in the same relevant conditions, and their expression can thus serve as a meaningful phenotypic readout.

Next, integrating RNA and protein measurements for joint analysis (**Methods**) highlighted gene programs that are either common across conditions, or unique to different conditions (**Figure 3I**), with the co-cultured cells uniquely enriched for induction of immune escape pathway genes and the ICR signature. Specifically, we learned programs in each treatment condition by an adapted jackstraw PCA procedure (**Methods**; **Figures S5A-S5G**), and annotated them by enrichment for functional gene categories (**Supplementary Data Table 5**). Several programs, including cell cycle regulation, DNA repair, and antigen presentation, were shared across conditions, whereas immune escape programs were uniquely recovered in co-culture data, and interferon response genes in IFNγ stimulation (**Figures 3I and S5H**; **Supplementary Data Table 5**). Thus, single-cell RNA and protein profiles provide rich relevant phenotypes with which to assess the impact of CRISPR perturbations.

### A computational model to infer perturbation effects from Perturb-CITE-seq

We developed a computational framework to model the effects of genetic perturbations on both RNA and protein profiles of individual genes across the cells in the screen (**Figure 4A; Methods**). Briefly, we used dial-out PCR data to determine the identity of perturbations (sgRNAs) in each cell (**Figure S6A; Methods**), and applied a linear model with elastic net regularization to infer the mean effect of each perturbation (sgRNA) on each feature (RNA and protein levels). We used a total of 4,461 RNA and 20 protein features, including all measured proteins and the union of the 1,000 top variable genes (**Methods**) and gene members of the programs identified by jackstraw PCA in any one condition. As we have previously shown (*2*), the detection of a sgRNA in a cell does not necessarily mean that this cell is perturbed, either because the sgRNA has not yet perturbed the gene, or the perturbation has not yet had an impact. To address this, we calculated the probability of a successful perturbation after fitting an initial regulatory matrix, and then updated sgRNA assignments using an estimate of the probability for each cell that it was successfully perturbed. Following our prior studies (*2*), we included both cell cycle and cell complexity (number of UMIs) as known covariates that impact cell profiles (**Figures 3F and S4C**). Including these covariates improved the model fit quality (residuals approach mean zero, independent and identically distributed; **Figures S6B and S6C**). Finally, we performed a permutation test to assess the empirical significance of each coefficient in the inferred regulatory matrix (**Methods**).

**Figure 4.**
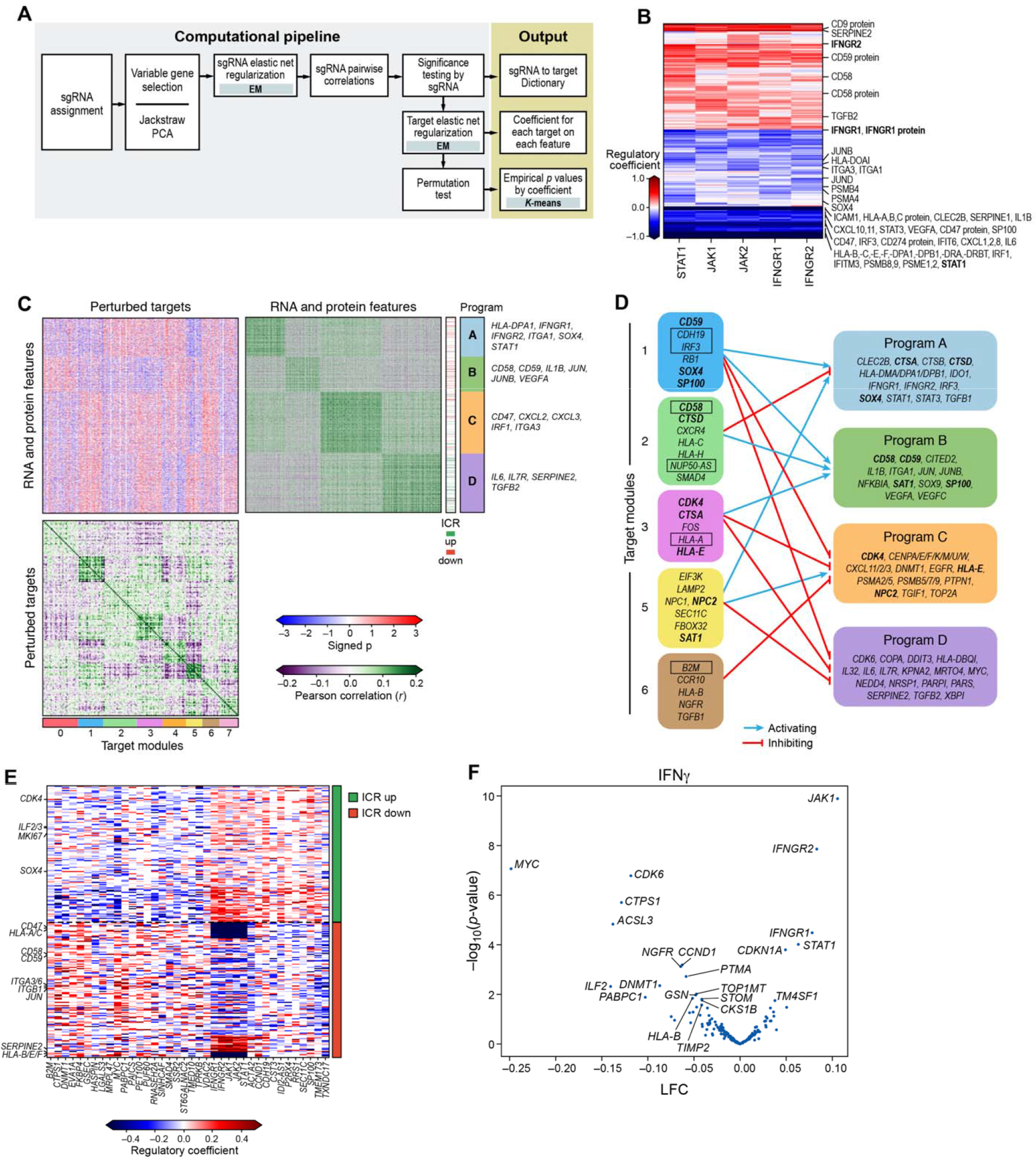
Perturb-CITE-seq reveals co-functional modules that are dependent- or independent of a predominant IFNg/jak-STAT mechanism. **(A)** Overview of computational approach (Methods). (B) Perturbations in JAK/STAT pathway affect known and putative mechanisms of immune evasion. Regulatory effect (values from the model shown in a; red/blue: perturbation induces/represses gene feature) on different RNA and protein features (rows; key genes are labeled) when perturbing different genes in the JAK-STAT pathway (columns). (C)(D) Co-functional modules and co-regulated programs in the Perturb-CITE-Seq screen. (C) Middle heatmap: Signed significance (– log_10_ (Empirical p) * sign(β), red/blue color bar) for the effect on each RNA or protein feature (rows) of perturbing each gene (columns, excluding JAK-STAT targets) in the co-culture condition. Right and bottom matrices: Pearson correlation coefficient (purple/green color bar) between the significance profiles of either gene/protein features (right matrix) or perturbed genes (bottom matrix). Co-functional modules (bottom bar) and co-regulated programs (right bar) are identified by *K*-means clustering of each of the bottom and right matrices separately (*k*=4 and 8, respectively), and the clustering defines the row and column order. (D) Schematic representation of the regulatory connections (blue/red: module gene activates/represses program) between select modules (left) and programs (right) from (C), highlighting key program genes. Bold font: select regulators that are also members of regulated programs. Boxed font: selected regulators significantly enriched/depleted in the viability screen (Figure 2D and 2E). (E)(F) The ICR program is coherently regulated by the perturbed regulators. (E) Regulatory effect (*β* values; red/blue: perturbation induces/represses gene feature) on different RNA and protein features from the ICR program (rows, sorted by genes in the induced as repressed ICR, as defined in Ref. 8) by perturbations of different genes in the screen (columns), clustered by *K*-means clustering (*K*=2). (F) Change in ICR signature scores (x axis, log_2_(fold-change) (LFC),) and its associated significance (y axis, −log_10_(P-value), Welch’s t test) for each perturbation (dot) in the IFNy condition (Methods). Key perturbations with significant effects are noted.

### Perturb-CITE-Seq model highlights established and novel functional modules that control diverse immune evasion pathways

The Perturb-CITE-Seq model correctly reconstructed the impact of genes known to affect resistance to immunotherapy, especially the effect of perturbing the IFNγ response machinery (**Figure 4B**), as well as novel pathways. The regulatory model can be parsed into eight major *cofunctional modules* of perturbed factors that similarly impact one or more of four major gene *coregulated programs* (**Figures 4C and 4D**). In particular, genes that were hits in the viability screen partitioned into different functional modules, thus highlighting the effect of the same (converging) or distinct (divergent) mechanism, which could not be distinguished by a viability screen. Those genes are also often members of regulated programs (**Figure 4D**).

First, the model accurately recovers the effect of perturbing components of the IFNγ response pathway, such that perturbation of any major known node of this pathway (*IFNGR1, IFNGR2, JAK1, JAK2, STAT1*) down-regulated a coherent RNA and protein regulatory program in coculture (**Figure 4B**). The downregulated program included key components of MHC Class I and Class II antigen presentation and associated machinery (*e.g., PSMB4, PSMB8, PSMB9, PSMA4, HLA-A,B,C* gene and protein, *HLA-E, HLA-F, HLAD-DPB1*), chemokines associated with antitumor immunity (*CXCL1, CXCL2, CXCL8, CXCL10, CXCL11*), inflammatory cytokines (*STAT3, IL1B, IL6*), interferon response elements (*STAT1, IRF1, IRF3, IFITM3, IFIT6*), surface checkpoints (*CD274 protein, CD47* gene and protein), and genes associated with cell differentiation states (*SOX4, ITGA3, ITGA1*). Thus, multiple features implicated in modifying the response to immunotherapies are directly regulated by the IFNγ-JAK/STAT axis, suggesting that some of these mechanisms are in fact a consequence of defective IFN’-JAK/STAT signaling, rather than independent modes of resistance.

Other transcripts and proteins were upregulated following perturbations in the IFN’-JAK/STAT nodes. These include *SERPINE2* and *TGFB2*, CD9 protein, CD59 protein, and both *CD58* transcript and CD58 protein (**Figure 4B**). It is likely that these induced genes are not regulated by the IFNγ-JAK/STAT pathway. For example, knockout or downregulation of *CD58* is associated with immune evasion (**Figures 2D, 2E, and 3D**), and thus its upregulation following perturbations in the IFNγ-JAK/STAT module is likely not part of this pathway’s immune evasion mechanism, and *CD58’s* role in immune evasion may be distinct from defects in the IFN’-JAK/STAT pathway.

To focus on other co-functional modules whose perturbation may affect distinct programs, we examined the regulation matrix after excluding perturbations in the IFNγ-JAK/STAT pathway genes (**Figure 4C**), and recovered novel regulators and mechanisms of immune evasion, either related to or distinct from the impact of the IFNγ-JAK/STAT pathway (**Figures 4C and 4D**). For example, *SOX4(9, 12), RB1(22), SP100(22) and IRF3(23*) all comprise one co-functional module (module 1), despite their different known roles in EMT (*SOX4*), cell cycle regulation (*RB1*), response to IFNγ (*SP100*) and transcriptional regulation of type I interferon response (*IRF3*) (**Figure 4D**). Interestingly, both SOX4 inactivation and RB1 inactivation (genetically or through hyperactivation of CDK4/6 phosphorylation) alter responses to immunotherapies in melanoma models (*12*) and breast cancer (*22*), respectively. Notably, *SOX4, IRF3* and *SP100* were repressed by IFNγ-JAK/STAT pathway in our screen (**Figure 4B**). The co-regulated program affected by their perturbation in our system (**Supplementary Data Table 6**) includes genes involved in cell cycle (*CDC80, CDKN3, CENPM*), metabolism (*COX20, DCTPP1, FH, IDO1, UCHL5, ZDHHC4, PAPSS1, PTGES*), mRNA maturation (*MRTO4, NSRP1, SNHG15*), DNA repair (*RAD51C*) and inflammation (*IFI16* and *IL1B*) or checkpoints (*IDO1*).

The perturbations also altered the expression of the ICR program, which we originally defined in patients (*2*), in both co-culture (**Figure 4E**) and IFN γ treatment (**Figure 4F**). Perturbations of the IFNγ-JAK/STAT module strongly increased the Overall Expression of the ICR signature (JAK1 p-value = 1.3e-10, IFNGR2 p-value = 1.4e-8, IFNGR1 p-value = 3.3e-5, STAT1 p-value = 9.6e-5, statistics by Welch’s T test, **Methods**), as did deletion of *TMEM173*, encoding STING (STimulator of INterferon Genes), which activates a type I interferon anti-tumor response (*24*) (**Figure 4E**). STING agonists are in clinical testing in melanoma patients with resistance to ICI therapy, and other cancers (*25*). In contrast, other perturbations, such as KO of *CDK6, MYC, ILF2, DNMT1*, or *ACSL3*, repressed the ICR signature (**Figure 4F**), consistent with our previously reported patient and pre-clinical data, where we demonstrate that upregulation of MYC and CDK4/6 (and their transcriptional targets) was associated with increased resistance to immunotherapy, and CDK4/6 inhibitors reduced resistance in human and a pre-clinical melanoma models (*9*).

Taken together, our Perturb-CITE-seq analysis rediscovered key mechanisms of immune evasion, organized them into modules and related them to the genes and programs they impact, as well as recovered new modules that may confer immune evasion beyond defects in the IFNγ-JAK/STAT pathway and antigen presentation.

### Loss or downregulation of CD58 confers immune resistance to T cell and NK cell mediated killing

From our integrated analysis across the viability and Perturb-CITE-seq data, *CD58* emerged as a compelling new factor in immune evasion: KO of *CD58* had a strong impact on resistance to T cell mediated killing in the viability screen and CD58 RNA/protein were down regulated in the co-culture system. Importantly, based on the Perturb-CITE-Seq screen, unlike many other genes needed for T cell mediated killing, *CD58* is not activated by the IFNγ-JAK/STAT pathway (**Figure 4B**), and the *CD58* KO conferred a different molecular phenotype than IFNγ-JAK/STAT pathway perturbations, belonged to a distinct module, and did not impact the expression of antigen presentation genes (**Figure 5A**). Thus, *CD58* loss may represent a resistance mechanism distinct from IFNγ-JAK/STAT pathway inactivation. CD58 is an adhesion protein typically expressed on the surface of antigen-presenting cells (APCs), where it binds to CD2 on CD8^+^ T cells and Natural Killer (NK) cells (*19*). Little is known about the potential role of CD58 in cancer, partly because there is no known mouse homolog to study it in pre-clinical models. Thus, patient-derived melanoma/TIL co-cultures provide a unique opportunity to study CD58 in the context of human immune evasion.

**Figure 5.**
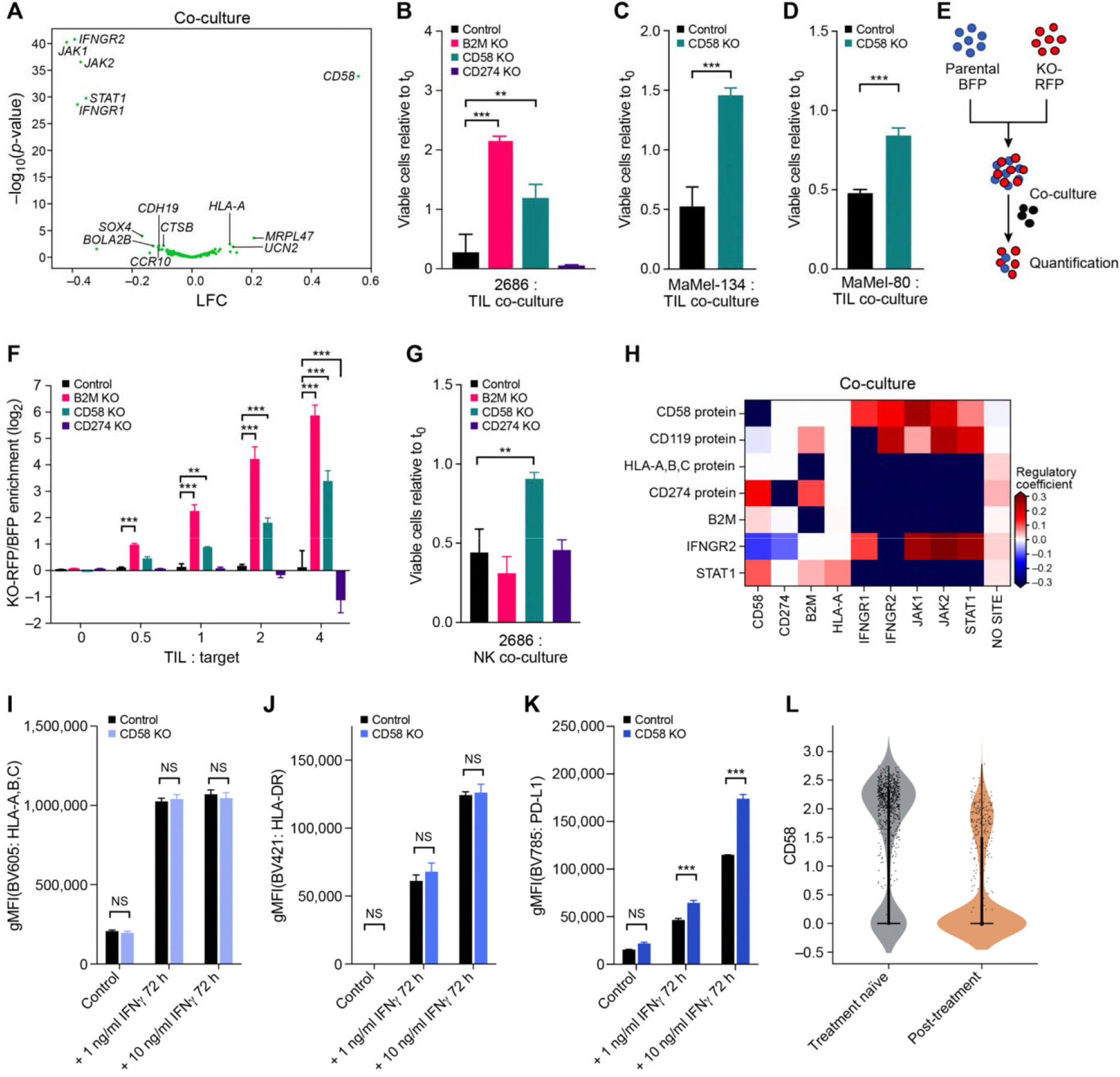
CD58 loss is a distinct mechanism of immune evasion from TIL and NK-cell mediated killing. **(A)** CD58 perturbation affects a distinct regulatory program. Change in signature scores of the CD58 regulatory program (x axis, log_2_(fold-change) (LFC),) and its associated significance **(y** axis, −log_10_(P-value), Welch’s t test) for each perturbation (dot) in the co-culture. Key perturbations with significant effects are noted. **(B)-(D)** CD58 KO enhances cell viability in TIL:cancer cell co-culture models. Ratio of viable cancer cells (y axis, relative to t_0_) in co-culture models of control (black), CD58 KO (turquoise), or B2M KO and CD274 KO (pink and purple, **(B)** only) cells (x axis) after 48h (b,d) or 72h **(C)** in co-culture. ns: p>0.05; * p<0.05; ** p<0.005; *** p<0.001; One-way ANOVA with Tukey *post hoc* test in b, t-test in c,d. Error bars: Mean ±SD. **(E)(F)** CD58KO enhances cell survival in a competition assay. **(E)** Competition assay schematic. BFP (blue) labeled parental cells and RFP (red) labeled KO cells are cocultured with autologous TILs and the ratio of RFP to BFP is calculated as an estimate of relative fitness (**Methods**). **(F)** RFP to BFP ratio (y axis, log_2_(RFP/BFP)) in a competition assay between BFP parental cells and RFP labeled cells with no perturbation (black), B2M KO (pink), CD58 KCO (turquoise), or CD274 KO (purple) after 48 hours of co-culture with different TIL ratios (x axis). ns, p>0.05; *, p<0.05; **, p<0.005; ***, p<0.001; 2-way ANOVA with Tukey *post hoc* test. Error bars: Mean ±SD. **(G)** CD58 KO confers resistance to NK cell mediated killing. Ratio of viable cancer cells (y axis, relative to t_0_) in an NK co-culture model of control (black), CD58 KO (turquise), B2M KO (pink), or CD274 KO (purple) cells (x axis) after 24h. ns: p>0.05; * p<0.05; ** p<0.005; *** p<0.001; One-way ANOVA with Tukey *post hoc* test. Error bars: Mean ±SD. **(H)-(K)** CD58 perturbation in co-culture does not affect B2M and HLA expression at the RNA and protein level but induces CD274. **(H)** Regulatory effect (*β* values from the model shown in a; red/blue: perturbation induces/represses gene feature) in Perturb-CITE-Seq on key RNA and protein (CITE) features (rows) when perturbing different genes in the JAK-STAT pathway, CD58 or CD247 (columns). **(I)-(K)** Surface expression (y axis) of MHC class-I (i), MHC class-II (j) or CD247 (k) at baseline and after stimulation with different levels of IFNγ for 72 hours (x axis), in control (black) and CD58 KO (blue) cells. ns: p>0.05; * p<0.05; ** p<0.005; *** p<0.001; one-way ANOVA with Tukey *post hoc* test Error bars: Mean ±SD. **(L)** CD58 transcript levels are higher in patients who failed immunotherapy treatment. Distribution of expression levels (y axis, log_2_(TPM+1)) of CD58 RNA in melanoma cells from tumors in patients who were either treatment naïve (grey) or were resected after failure of immunotherapy (tan) in the scRNA-seq data from the ICR-signature discovery cohort(*9*).

To further validate our finding that loss or downregulation of CD58 causes impaired T cell mediated toxicity, we knocked out each of *CD58, B2M* or *PD-L1* by Cas9/sgRNA ribonucleoprotein electroporation in each of three patient-derived melanoma cell lines and cocultured them with their autologous TILs (**Figures S7A-S7C; Methods**). In all three independent models, *CD58* KO and *B2M* KO conferred resistance to T cell mediated toxicity (**Figures 5B-5D**), whereas *PD-L1* KO sensitized cells and resulted in higher T cell mediated killing (**Figures 5B**). The different KO did not impact proliferation rate (**Figure S7D**), caspase-mediated apoptotic potential, or sensitivity to chemotherapy with dacarbazine (**Figure S7E**), showing that increased survival was specifically due to escape from T cell mediated killing rather than non-selective differences in survival of different genotypes. Next, we performed competition assays, where BFP-expressing parental and RFP-expressing KO cell lines were mixed at a 1:1 ratio and co-cultured with TILs in the same well (**Figure 5E**). Both *CD58* KO and *B2M* KO cells had a strong competitive advantage over parental cell lines and PD-L1 KO, in a T cell dose-dependent manner, with the *B2M* KO advantage being more pronounced (**Figure 5F**). Taken together, these experiments validate our screen results that *CD58* KO confers immune evasion to antigen-targeted T cell killing.

Because CD2, the CD58 binding partner, is also expressed on Natural Killer (NK) cells, we hypothesized that *CD58* KO would also increase immune evasion from these cells. To test this hypothesis, we co-cultured melanoma cells with *CD58, B2M* or *PD-L1* KO as well as unperturbed control cells, with human NK cells (NK-92, **Methods**). Supporting our hypothesis, *CD58* KO conferred immune evasion. Because NK cells cause tumor lysis independent of MHC Class I expression, *B2M* KO cells were lysed at similar rates as parental and *PD-L1* KO (**Figure 5G**), while *CD58* KO were resistant to NK killing.

### CD58 confers immune evasion without affecting antigen-presentation, and is not induced by IFNγ

Because *CD58 KO* conferred immune evasion from both T and NK cells, we hypothesized that its mechanism of action was independent of antigen presentation via MHC. Notably, in our Perturb-CITE-seq data *CD58* KO did not significantly alter the level of the *B2M* transcript or MHC Class I protein (encoded by *HLA-A*) (**Figure 5H**). To further validate this, we performed flow-cytometry for MHC Class I (**Figure 5I**) and II (**Figure 5J**) of *CD58* KO *vs.* isogenic parental cells at baseline and following stimulation with low-dose or high-dose IFNγ. Indeed, there were no significant differences in MHC expression between parental and *CD58* KO cells, demonstrating that this perturbation does not alter baseline stability or the IFNγ-induction of MHC proteins. Our Perturb-CITE-seq data also suggested that *CD58* KO led to increased expression of PD-L1 (**Figure 5H**). Using flow-cytometry, we confirmed that stimulation with low- or high-dose IFNγ resulted in higher levels of PD-L1 protein in *CD58* KO cell lines compared to parental control (**Figure 5K**). This raises the possibility that upregulation of PD-L1 could contribute to immune evasion in *CD58* KO in T cell co-culture. Notably, *CD58* KO impacts the expression of additional genes (**Figures 4C, 4D, and 5H**), including a possible regulation of IFNGR2, which can provide additional contributions to its overall impact.

Conversely, neither B2M nor HLA-A KO impacted CD58 protein levels in our Perturb-CITE-Seq screen (**Figure 5H**), while *B2M* KO abrogated MHC Class I protein as expected and validated by flow-cytometry (**Figure S7A**). Moreover, perturbations of the IFNγ-JAK/STAT nodes (*JAK1, JAK2, STAT1, IFNGR1, IFNGR2*) actually led to an increase (not a decrease) in CD58 RNA and protein (**Figures 4B and 5H**), and stimulation with IFNγ (at 1 ng or 10 ng/mL) did not increase protein abundance of CD58 (PD-L1 was strongly induced at either dose, as expected) (**Figures S7F-S7I and 3B**), suggesting that CD58 is not induced via the IFNγ pathway.

Finally, we determined the level of mRNA expression of *CD58* in melanoma patients with resistance to ICI. Compared to treatment-naïve patients, ICI resistant patients had a significantly lower expression of *CD58* (**Figure 5L**). In line with our other data, this suggests that *CD58* downregulation (or loss) is associated with immune evasion from T cell mediated toxicity and resistance to immune checkpoint inhibitors in melanoma patients.

## DISCUSSION

In this study, we developed Perturb-CITE-Seq, a pooled CRISPR-Cas9 screen with multi-modal single-cell profiling readout, and used it in patient-derived tumor-immune models to dissect cancer cell mediated modulators of T cell mediated killing. We used a broadly applicable computational and statistical framework for integrated analyses of such screens, which accounts for key co-variates, including cell quality and cell cycle status, focuses on cells harboring impactful perturbations, generates a detailed regulatory model, and partitions it to interpretable co-functional modules that control co-regulated programs.

Our rich, multi-modal screen is critical to dissect which genes are part of a shared mechanism of resistance, which represent distinct mechanisms, and through which gene programs each act. First, Perturb-CITE-seq data readily identified major known clinical mechanisms of immune evasion, especially perturbations and their associated cell programs in the IFNγ pathway and downregulation or defects of the antigen-presentation axis. Perturbations of different nodes within IFNγ pathway led to highly converging molecular phenotypes, irrespective of the level of the defect. This is consistent with genomic profiling in patients with resistance to either anti-CTLA-4 (*26*) or anti-PD-1 therapy (*7*), and further emphasizes the role of IFNγ in T cell mediated anti-tumor immunity. Some of the genes down-regulated by IFNγ pathway perturbations, such as *CXCL10* and *CXCL11*, are genes whose genetic loss or downregulation was previously associated with immune evasion, suggesting that these do not represent a separate mechanisms of immune evasion. In contrast, other genes whose perturbation enhanced immune evasion in our screen (*e.g., CD58, CD59*) appear to reflect distinct mechanism, both because their expression was not impaired by IFNγ signaling defects, and because the molecular phenotype following their perturbation is distinct. Because many gross phenotypes in biology (immune evasion, cell cycle, viability, differentiation, etc.) are affected by multiple pathways and involve interactions between cells, this approach should provide a powerful solution in many other systems.

In particular, our Perturb-CITE-Seq and viability screens highlighted *CD58*, as a gene whose knockout enhances resistance to T cell mediated killing, a member of a co-functional module with a distinct phenotype, and a target whose RNA and protein levels are reduced in co-culture, but are not activated by the IFN γ pathway. Because there is no known mouse homolog of *CD58*, this target was not discovered in previously reported CRISPR screens performed in mouse models (*11–13*). While it was a top ranking hit in an immunotherapy screen in a human-engineered system (*10*), its role remained poorly understood. In patients with diffuse-large B cell lymphoma (DLBCL) (*27*), *CD58* mutations were concurrent with mutations in *B2M* leading with loss of antigen presentation. Because the two mutations occurred concurrently, it remained unclear whether these are independent mechanisms of immune escape. Our experiments and analysis show that loss of *CD58* confers immune evasion to a similar extent as loss of MHC Class I expression itself (through *B2M* KO), but through an independent path, and without impacting the expression of antigen presentation genes and proteins. Furthermore, PD-L1 is upregulated in *CD58* KO, suggesting that *CD58* loss could confer immune evasion directly (reduced adhesion) or indirectly (inhibitory PD-L1). *CD58* KO also impacted NK-mediated killing (**Figure 5G**), which may have implications for NK cell-based therapies in tumors with loss of MHC Class I presentation. Notably, the CD58/CD2 axis is a potent co-stimulatory pathway in CD8^+^ T cells that lack expression of CD28 (CD8+CD28-T cells) (*28*), which are common in the tumor-microenvironment and peripheral blood of solid tumor patients (*29*), and were the predominant T cells in our Perturb-CITE-seq screen (**Figure S4B**). Therapeutic engagement of the CD58/CD2 axis, either through stabilization of CD58 on the membrane or by stimulation of CD2, may represent a therapeutic opportunity, for example by manipulation of regulators of CD58 RNA and protein expression.

Our study is the first large-scale CRISPR-Cas9 screen with multi-modal single-cell readouts, providing a general approach, as well as addressing a critical clinical challenge in a unique patient-derived cell culture model system. We recover the landscape of resistance mechanisms to immunotherapies and, guided by our high-dimensional data, validate a novel mechanism of immune resistance. Large-scale multi-modal screens, spanning RNA, proteins, chromatin accessibility and imaging features, should enable unprecedented discovery across diverse biological systems, and detailed dissection of complex cellular and inter-cellular circuits.

## Acknowledgements

The Koch Institute-Dana-Farber/Harvard Cancer Center Bridge Project Grant (Av.R., B.E.J., B.I.), the Klarman Cell Observatory (Av.R., O.R.R.), HHMI (Av.R.), NHGRI Center of Excellence in Genome Science (CEGS) the Center for Cell Circuits (Av.R.). Av.R was an Investigator of the Howard Hughes Medical Institute. K08CA222663 (B.I.), U54CA225088 (B.I.), Burroughs Wellcome Fund Career Award for Medical Scientists (B.I.), Louis V. Gerstner, Jr. Scholars Program (B.I.), Ludwig Center for Cancer Research at Harvard (K.W.), NIH F32 fellowship F32AI138458 (P.T.), the Broad Fellows Program (B.C.). This research was funded in part through the NIH/NCI Cancer Center Support Grant P30CA013696 at Columbia University.

## Data availability

Processed data will be available via the single-cell portal upon publication in a peer-reviewed journal: https://singlecell.broadinstitute.org/single_cell/study/SCP1064/perturb-cite-seq-pooled-screens-in-patient-models-identify-loss-of-cd58-as-novel-mechanism-of-cancer-immune-evasion#study-summary. Upon publication in a peer-reviewed journal, raw data will be available on the Broad Data Use and Oversight System https://duos.broadinstitute.org.

## Code availability

Code will be available on KCO GitHub.

## METHODS

### Patient derived melanoma cell culture

Patient derived melanoma cell lines were grown on non-pyrogenic, polystyrene tissue culture treated plastic ware (Corning, Corning, NY) in Roswell Park Memorial Institute (RPMI) 1640 Medium supplemented with 10% heat-inactivated fetal bovine serum (FBS), GlutaMax, 10 mM HEPES, 10 mg/L Insulin, 5.5 mg/L Transferrin, 6.7 μg/L Sodium Selenite, and 55 μM 2-Mercaptoethanol (all Thermo Fisher Scientific, Waltham, MA) at 37°C and 5% CO_2_ in a humidified incubator. Melanoma cell line 2686 and matched tumor infiltrating lymphocytes (TILs) were previously derived under IRB approved protocols and provided by MDACC, Texas, USA (*9, 14*). Melanoma cell lines Ma-Mel-80 and Ma-Mel-134 and corresponding TILs were derived un IRB approved protocols and provided by UK-Essen, Germany. Melanoma cells tested repeatedly negative for contamination with mycoplasma and other contaminants using PlasmoTest (InvivoGen, San Diego, CA).

### Rapid T cell expansion from tumor infiltrating lymphocytes

Previously established TILs (*14*) were grown in 24 well plates in RPMI 1640 supplemented with heat-inactivated 10% human AB serum (Corning, Corning, NY), GlutaMax, 10 mM HEPES, 55 μM 2-Mercaptoethanol (all ThermoFisher Scientific, Waltham, MA), 50 Units/mL penicillin, and 50 μg/mL streptomycin at 37°C and 5% CO_2_ in a humidified incubator. TIL media was supplemented with 300-3000 IU recombinant human IL2 (Chiron Therapeutics, Emeryville, CA) and media with IL2 was replenished every two days by half media exchange. For rapid expansion, 0.5-1×10^6^ proliferating TILs were seeded with 100×10^6^ normal donor peripheral blood mononuclear cells irradiated with 5,000 rad in a 1:1 mixture of TIL media and AIM-2 media (ThermoFisher), supplemented with 30 ng/mL OKT3 (Miltenyi Biotec, Bergisch Gladbach, Germany) in a G-Rex-10 bottle (Wilson Wolf, St. Paul, MN). On day 2, 6,000 IU/mL IL2 were added followed by a half media exchange on day 5 and addition of another 6,000 IU/mL IL2. On day 7, cells were counted and a minimum of 6×10^7^ cells were transferred to a G-Rex-100 bottle in a total volume of 600 mL AIM-5 medium supplemented with 6,000 IU/ml IL2. On day 10, 400 mL AIM-5 with 6,000 IU/ml IL2 were added. On day 12, 3,000 IU/ml IL2 were added. Expansion was completed on day 14 and cells were frozen in aliquots of 10-100×10^6^ in Bambanker freezing medium (Nippon Genetics, Tokyo, Japan)

### Tumor infiltrating lymphocyte characterization

Expanded TILs were thawed and rested in TIL media supplemented with 3,000 U/mL rhIL2. On day three, 1×10^5^ TILs were seeded per well of a round bottom 96 well plate and stimulated with phorbol-12-myristate-13-acetate/Ionomycin in the presence of Brefeldin A (Biolegend) for 4 hours as per manufacturer’s instruction. Cells were washed once in cold PBS and stained for 15 minutes with Zombi NIR (Biolegend) diluted 1:500 in PBS to label dead cells. Cells were then washed once with full staining buffer (3% FBS in PBS with 2 mM EDTA) and surface antigens were stained for 30 min on ice using the following antibodies: gdTCR-BV421 (B1, Biolegend, 331218), CD45-Pacific Blue (HI30, Biolegend, 304022), CD3-AF532 (eBiosciences, 58/0038-42). Nen, cells were washed twice in ice cold staining Buffer and fixed and permeabilized using the eBioscience™ FoxP3/Transcription Factor Staining Buffer Set (ThermoFisher) per manufacturer’s recommendation. Intracellular antigens and fixation-stable surface antigens that might be internalized during activation were stained for 45 minutes on ice using the following antibodies: CD8a-BV570 (RPA-T8, Biolegend, 301038), Granzyme B-PE-CF594 (GB11, BD, 562462), FoxP3-APC (236A/E7, eBiosciences, 17-4777-42), CD4-AF700 (RPA-T4, Biolegend, 4345826), IFNγ-AF488 (4S.B3, Biolegend, 502517), IL-17A-PE-Cy7 (BL168, Biolegend, 512315), and TNF-a-APC-Cy7 (Mab11, Biolegend, 502943). Fluorescent-minus-one (FMO) staining controls were prepared for gating of IFN-g, TNF-a, IL-17A, and Granzyme B. After staining, cells were washed twice with permeabilization buffer, resuspended in staining buffer, and analyzed on a Sony SP6800 spectral analyzer (Sony, Tokyo, Japan)

### Lentivirus production for fluorescent protein expression

Lentivirus encoding nucleoplasmin nuclear localization sequence (NLS) tagged fluorescent proteins dsRed (NLS-dsRed) and blue fluorescent protein (NLS-BFP) were generated using pHAGE vectors (Gift from W. Marasco). To generate lentivirus, HEK293T cells (ATCC) were seeded in 6-well plates in DMEM supplemented with 10% FBS (D10). The next day, cells had reached 80% confluence and were transfected with plasmids for lentivirus production using TransIT-LT1 transfection reagent (Mirus Bio, Madison, WI). 3 μl transfection reagent was added to a total volume of 15 μL Opti-MEM (ThermoFisher) and incubated for 5 minutes at room temperature. In the meantime, the plasmid mix was prepared by mixing 500 ng packaging plasmid psPAX2 (addgene #12260, gift from Didier Trono) and 250 ng envelope plasmid pMD2.G (addgene #12259, gift from Didier Trono) with 500 ng pHage expression vector in 37.5 μL Opti-MEM. The transfection reagent mix was added to the plasmid mix, incubated for 30 minutes at room temperature and then added dropwise to the cells. All volumes reported are for the transfection of one well and were scaled for transfection of multiple wells and plates. Cells were incubated for 18 hours, and then the transfection media was removed and DMEM supplemented with 20% FBS (D20) was added. Supernatant containing lentivirus was collected after 24 hours and fresh D20 was added. After another 24 hours, media was again collected and the lentivirus containing media was pooled, filtered through a 0.45 μM syringe filter (ThermoFisher) to remove debris, aliquoted, frozen and stored at −80°C.

### Transduction of NLS-fluorescent proteins

Human melanoma cells were seeded in 12 well plates with increasing volumes of lentiviral supernatant in a final volume of 2 mL with 4 μg/mL polybrene (Millipore, Burlington, MA). Plates were centrifuged for 2 hours at 1,000x g at 30°C and cells were incubated for 16 hours at 37°C and 5% CO_2_ in a humidified incubator. The next day, cells were detached and seeded in 6-well plates. Cell lines with a transduction efficiency of 30% were sorted on a FACS Aria cell sorter, expanded and stored frozen in Bambanker (Nippon Genetics) until further use.

### Miniaturized imaging-based co-culture assay

10^4^ melanoma target cells expressing NLS-dsRed were seeded per well of a black walled 96-well plate (Corning) and treated for 16h with 1 ng/mL recombinant human Interferon gamma (Peprotech). After 16h, media were replaced with 100 μL full melanoma media without phenol-red containing 4μM Caspase-3/7 activity dye (CellEvent, ThermoFisher). Plates where imaged using a Celigo Imaging Cytometer (Nexcelom, Lawrence, MA) to obtain time point 0 cell counts. TILs were thawed 3 days prior to co-culture and cultured in TIL media with 3,000 IU/IL2 or re-activated on plates coated with 100ng/ml OKT3 in PBS when comparing preactivated TILs to rested TILs. On the day of the assay, TILs were collected, centrifuged and resuspended in TIL media without IL2, counted and increasing ratios of TILs were added to the melanoma target cells to a complete volume of 200 μL per well (final Caspase activity dye = 2 μM) and the co-cultures were incubated at 37°C and 5% CO_2_ in a humidified incubator. The plates were reimaged 24, 48 and 72h after TIL addition. Viable target cells were counted using Celigo Imaging Cytometer software. First, cells were identified by size and intensity of the NLS-fluorescent protein. Debris was defined by adjusting gating parameters to exclude condensed, late apoptotic cells and Caspase3/7-dye intensity was measured across all cells in the green channel to identify apoptotic cells. Finally, viable cells were defined as *viable Cells (Class) = NLS-dsRed positive AND NOT debris size AND NOT Caspase3/7 positive.* Viable target cell counts were normalized to the respective well counts on time = 0 using Excel (Microsoft, Redmond, WA) and plotted using GraphPad Prism 8 (GraphPad, San Diego, CA).

### Cas9 lentivirus production

Cas9 lentivirus was generated as outlined for other vectors above using 6 well plates and the expression vector pLX311-Cas9 (addgene #118018, gift from William Hahn and David Root),

### Lentiviral Cas9 transduction of human melanoma cell lines

Human melanoma cells were seeded in 12 well plates with increasing volumes of lentiviral supernatant in a final volume of 2 mL with 4 μg/mL polybrene (Millipore, Burlington, MA). Plates were centrifuged for 2 hours at 1,000x g at 30°C and cells were incubated for 16 hours at 37°C and 5% CO_2_ in a humidified incubator. The next day, cells were detached and seeded in 6-well plates. In parallel, 1×10^5^ cells of infected and non-infected cells were seeded in 12 well plates to be used for transduction efficiency assessment. On day 2 after infection, selection was initiated with 2 μg/mL Blasticidin S (ThermoFisher). When all cells in the non-infected but selected control were dead, transduction efficiency was calculated as a ratio of cells surviving selection divided by cells in the non-selected control. Cells with approximately 30% transduction efficiency were chosen for further experiments.

### Cas9 activity assay

To assess Cas9 editing activity, Cas9 expressing human melanoma cell line 2686 was transduced with lentiviral particles produced as described above using expression plasmid pXPR_011 (Addgene #59702, gift from John Doench and David Root), encoding for Enhanced Green Fluorescent Protein (EGFP) and a guide RNA targeting EGFP (*31*). Cells with no Cas9 expression were transduced as non-editing, EGFP positive controls. Transduced cells were selected in media supplemented with 0.75 μg/mL puromycin. After selection was completed, EGFP expression was assessed by flow cytometry on a Sony SP6800 spectral analyzer (Sony, Tokyo, Japan). Control cells with and without EGFP expression were used as gating controls and Cas9 activity was defined as %EGFP negative cells.

### Generation of pooled Perturb-CITE-seq gDNA library

To generate a Perturb-CITE-Seq vector compatible with FACS selection, we modified the lentivirus CROPseq-Guide-Puro vector (*4*) by Gibson Assembly to insert a gene expressing the fluorescent protein mKate2 (CROPseq-mKate2). After SPRI-purifying CROPseq-Guide-Puro which had been digested with BsiWI, we performed a Gibson Assembly to insert the Kate2 expressing oligo ordered as a gene string from Invitrogen (**Supplementary Data Table 7**).

For the pooled Perturb-CITE-seq CRISPR library, we designed a 744 single-guide (sg)RNA gDNA library targeting 248 genes selected from a previously defined signature of immune checkpoint inhibitor resistance in melanoma (*9*), with three gRNAs targeting each gene with sequences picked from the Broad Institute Genetic Perturbation Platform Web sgRNA Designer (https://portals.broadinstitute.org/gpp/public/analysis-tools/sgrna-design) (*32*). We also incorporated two types of control gRNAs, non-targeting and non-gene targeting intergenic controls, in the library, each at 5% of the total guide count (n=74 control guides for Perturb-CITE-Seq experiment). In addition to this library for Perturb-CITE-Seq, we generated a separate library (extended control library), including the same targeting sgRNAs, but a larger number of control guide RNAs (n=758,) for the large-scale CRISPR viability screen to increase the power for enrichment/depletion detection. All guide sequences can be found in Supplementary Data Table 1. The pool of guide sequences with appropriate overhangs for Golden Gate cloning was synthesized by Twist Biosciences. The gRNA oligo pool was amplified by PCR and inserted into a BsmBI-linearized CROPseq-mKate2 vector backbone via a Golden Gate ligation reaction. The resulting product was purified by isopropanol precipitation prior to transformation into electrocompetent Lucigen Endura cells (10 μF, 600 Ohms, 1800 V). Transformed libraries were grown overnight in shaking liquid culture at 37°C, and a library representation of >200 colonies per guide was determined prior to CRISPR library pool plasmid purification by Endofree Maxi (Qiagen). Coverage of the 818 guide library for Perturb-CITE-Seq as well as the extended control library was confirmed by PCR amplification of the guide sequence from the final library pool and deep sequencing on an Illumina MiSeq.

### Perturb-CITE-seq ICR gDNA library lentivirus production

Lentivirus of the Perturb-CITE-seq sgDNA library was produced in multiple 6-well plates as described for other vectors above. To generate the extended control ICR library, the ICR library was mixed at equimolar ratios with the extended control library prior to transfection of the HEK cells.

### ICR-library titration

2.5×10^5^ Cas9-expressing melanoma target cells (2686) were seeded in 1 ml of media in each well of a 12-well plate in the presence of 4 μg/ml polybrene (Millipore). ICR-library virus was added at increasing dilution with one well receiving no virus and serving as non-transduction control. The plates were centrifuged at 1000 x g for 2 h at 30°C and incubated overnight at 37°C and 5% CO_2_ in a humidified incubator. After 16h, viral supernatants were removed, and fresh media was added. 8 hours later the cells were split in two plates and selection with 0.75 μg/ml puromycin was initiated in one of the plates. When all cells in the non-infected but selected control were dead, transduction efficiency was calculated as a ratio of cells surviving selection divided by cells in the non-selected condition. At the same time transduction efficiency in the non-selected cells was assessed by detecting the fluorescent marker mKate2 by flow cytometry using a Sony SP6800. Multiplicity of infection (MOI) was calculated as MOI = LN(1-x/100) with x=%mKate^+^ or x=%survivors respectively. The percentage of cells receiving exactly one viral particle was calculated as Single-Infection%=((1-x/100)*LN(1-x/100))/(x/100)*100 as previously described (*12*).

### Introducing ICR-libraries in melanoma target cells

Cas9-expressing melanoma target cells (2686) were transduced with the ICR-library (818 sgRNAs, for scRNA-seq) or the ICR-extended library (818 sgRNA from the ICR library + 818 sgRNA controls, for enrichment screening) aiming for a transduction rate <10% (>90% single infection rate) and at least 1000x coverage per sgRNA. To this end, 2.5×10^6^ cells per well of a 12 well plate were seeded in the presence of 4 μg/ml polybrene and titrated ICR-library virus was added. Cells with virus were centrifuged at 1000 x g for 2 h at 30°C and incubated overnight at 37°C and 5% CO_2_ in a humidified incubator. After 16h, viral supernatants were removed, cells were detached and reseeded in T175 cell culture flasks, and Puromycin selection was started 24h after transduction. A 6-well plate with and without selection plated with transduced and nontransduced cells served as transduction control. Selection was completed after day 6 and transduced cells were further propagated in the presence of puromycin. On day 7 and day 14, aliquots of 1×10^6^ cells were saved for gDNA read-outs.

### Large-scale co-culture assay for CRISPR-Cas9 viability screen

Melanoma cells were transduced with an ICR-library extended with an equal number of control-sgRNAs to increase power for enrichment detection, as described in library preparation above, and three pellets of 1×10^6^ perturbed target cells were collected on day 7 to later quantify drop-out of essential genes. After 14 days, perturbed melanoma cells were plated in 12 well plates with 1.25×10^5^ cells/well (~900 x guide representation per plate). At the same time, three aliquots of 1×10^6^ melanoma cells were collected to quantify the distribution of sgDNAs at the beginning of the experiment. Cells were treated with 1 ng/ml recombinant human IFNγ (in both treatment and co-culture conditions) or left untreated and incubated at 37°C and 5% CO_2_ in a humidified incubator. After 16 hours, media were replaced with fresh pre-warmed media without IFNγ. TILs had been thawed three days prior to the experiment and cultured in TIL media supplemented with 3,000 IU/ml rhIL2. On the day of TIL addition, TILs were resuspended, pelleted at 400 x g at 4°C for 5 minutes, resuspended in TIL media without IL2, counted, and added at effector to target ratio ranging from 1:1 to 4:1 to the perturbed IFNγ pretreated melanoma target cells in a final volume of 2 ml. Plates were centrifuged at 400 x g for 5 minutes to ensure cell-to-cell contact and the co-cultures were kept for 48 h at 37°C and 5% CO_2_ in a humidified incubator. For the untreated control and IFNγ treatment the experiment was set up in triplicate plates. For each of the co-culture ratios one plate was set up. After 48 hours, each plate was washed once with ice cold PBS to remove TILs and dead cells, surviving cells were detatched using accutase, pelleted, and pellets were stored at −80°C until further processing. Surviving target cells on distinct plates containing all conditions as triplicates were counted to quantify target cell killing to verify accurate immune selection pressure as previously determined in the miniaturized coculture experiments (above). Genomic DNA was extracted and purified from cells using QIAamp DNA Micro Kit (Qiagen 56304) with the user-developed protocol QA43 and no more than ~0.5M cells per column. sgDNA sequences were amplified from genomic DNA for Illumina sequencing in two PCR steps. The first PCR was conducted in parallel 50 μL reactions, ensuring no more than 500ng of genomic DNA template per reaction. Using primers 642F and 643R and NEBNext High-Fidelity 2X PCR Master Mix (NEB M0541L), PCR was run for 10 cycles with the following parameters: 98°C denaturation for 60 sec, 68°C annealing for 30 sec, and 72°C extension for 60 sec. Taking 5 μL of from PCR1 for another 50 μL reaction, using primers 997 and 998 to add Illumina adapters and NEBNext master mix, PCR2 was run for 10 cycles with the following parameters: 98°C denaturation for 15 sec, 62°C annealing for 15 sec, and 72°C extension for 16 sec. With 5μL of PCR2 product, qPCR was conducted with SYBR green and the same primers and parameters as PCR2 to estimate the number of PCR cycles to reach sufficient amplification for Illumina sequencing. With the same primers and parameters as PCR2, additional cycles were run for each reaction as calculated. PCR products were then purified by 1X SPRI and prepared for sequencing on an Illumina HiSeq 2500 instrument in RapidRun mode with custom read 1 (primer 503F). All primer sequences are listed in **Supplementary Data Table 8**.

### Viability screen analysis

CRISPR/Cas9 screens from genomic DNA were analyzed using MAGeCK and MAGeCKFlute software packages (*33*). Briefly, MAGeCK maps sequencing reads to the reference library of sgDNA and returns the quantity of reads that confidently mapped to each sgDNA sequence without allowing for mismatches. To control for sequencing depth, sgDNA counts are median-normalized. Several metrics were used to evaluate the quality of each sequenced sample, including Gini index, missed gDNAs, and correlation of sgDNA counts between replicates. MAGeCK’s robust rank aggregation (RRA) method was used to discover significantly enriched or depleted genes in test versus control conditions.

### Perturb-CITE-seq co-culture experiment

After 14 days of editing and selection, 1.25×10^5^ Cas9-expressing, ICR-library perturbed melanoma target cells were seeded per well of a 12-well plate (One Plate with 1.5×10^6^ cells total per condition, >1,800x representation). Cells were treated with 1 ng/ml recombinant human IFNγ (in both treatment and co-culture conditions) or left untreated and incubated at 37°C and 5% CO_2_ in a humidified incubator. After 16 hours, media were replaced with fresh pre-warmed media for the co-culture group and cells of the control and treatment groups were washed once with cold PBS, detached using Accutase (Stem Cell Technologies, Vancouver, Canada), quenched with full cold media, pelleted at 400 x g at 4°C for 5 minutes, resuspended in CITE staining buffer (PBS with 2% BSA and 0.01% Tween-20), filtered through a pre-wetted 70 μM cell strainer (Corning), counted, and 1×10^6^ cells were aliquoted in 1.5 ml DNA LoBind microcentrifuge tubes (Eppendorf, Hamburg, Germany) and pelleted again. Cells were resuspended in 100 μL CITE staining buffer, and 5 μL TruStain FcX blocking antibodies (Biolegend) were added and incubated for 10 minutes on ice. Cells were again pelleted and resuspended in 100μL CITE staining buffer and 5μL of CITE-seq antibody pool (~1:500 final dilution for each antibody) were added to each sample and incubated for 30 min on ice. Nexy, cells were washed for a total of 3 washes with 100μL CITE staining buffer and then processed for scRNA-Seq with 15,000 cells were loaded onto each of 8 channels per condition using the 10x Chromium system with the Chromium Single Cell 3’ Library and Gel Bead kit v3 (10X genomics, Pleasanton, CA) per manufacturer’s instructions.

For the co-culture condition, TILs had been thawed three days prior to the experiment and cultured in TIL media supplemented with 3,000 IU/ml rhIL2. On the day of TIL addition, TILs were resuspended, pelleted at 400 x g at 4°C for 5 minutes, resuspended in TIL media without IL2, counted, and 2.5×10^5^ cells were added in a final volume of 2 ml to the perturbed IFNγ pretreated melanoma target cells (final effector/target ratio = 2:1). Plates were centrifuged at 400 x g for 5 minutes to ensure cell-to-cell contact and the co-cultures were kept for 48 h at 37°C and 5% CO_2_ in a humidified incubator. After 48 hours, media was removed from the co-culture plates and the cells were washed once with cold PBS and detached as described above. After initial centrifugation, cells were resuspended in ice-cold PBS with 1:500 dilution of Zombi-NIR dead cell stain (Biolegend) and incubated for 15 minutes in the dark on ice. Cells were then washed once with CITE staining buffer, resuspended, filtered, counted, aliquoted and Fc-blocked as described above. CD45-Pacific Blue (HI30, Biolegend, 304022) was included during the CITE antibody staining to label TILs for sorting. After three washes in CITE buffer, cells were again filtered and viable melanoma target cells were sorted using a FACS Aria III with cooling system. Melanoma target cells were identified using cell gating with FSC-A and SSC-A, doublet exclusion using FSC-A and FSC-H, gating on Zombi-NIR dim cells and then gating on CD45 negative, mKATE2 positive target cells. After sorting, cells were centrifuged once at 400 x g for 5 minutes at 4°C, resuspended in CITE staining buffer and 15,000 cells/channel were loaded onto the 10x Chromium system as described above.

### Perturb-CITE-Seq library generation and sequencing

After loading of cells onto the 10X Chromium system (Chromium Single Cell 3’ Library and Gel Bead kit v3) single cell gene expression libraries were generated following the manufacturer’s instructions with one change during cDNA amplification: to increase abundance of CITE-seq oligos, 2 pmol of the ADT additive was spiked-in into the 10X cDNA amplification. Following cDNA amplification, a 0.6X SPRI was performed to separate large gene expression library cDNA from smaller CITE-seq oligos. The supernatant of this 0.6X SPRI was set aside to generate the CITE-seq expression library. The 0.6X cDNA amplification SPRI was then completed and gene expression libraries were generated following the manufacturer’s instructions. Returning to the supernatant of the cDNA amplification 0.6X SPRI, 1.4X SPRI was added to the initial 0.6X SPRI supernatant. This double sided SPRI was completed following two 200 μL 80% ethanol washes. The product was eluted in 50 μL TE buffer, and a second 2.0X SPRI was performed on this CITE-seq template eluting in 50 μL TE buffer. To generate the CITE-seq sequencing library, 5 μL of the SPRI-cleaned CITE-seq template was mixed with 25 μL of 2X NEBNext HiFi Master Mix, 11.5 μL of diH2O, and 1 μL of a 1:1 ADT primer mixture (10 μM stock concentration of each primer). To enable multiplexing, one of six unique P7 indices were included in each reaction. The CITE-seq sequencing library reaction was cycled with the following conditions using a thermocycler: 98°C for 10 minutes, (98°C for 2 sec, 72°C for 15sec)x18 then 72°C for 1 minute.

To amplify fragments containing the guide barcodes, a 10 ng fraction of the WTA was mixed with 15 μL Phusion High-Fidelity PCR Master Mix, 11.5 μL H_2_O, 1.25 μL CROPDialOut_R1 (25 μM), and 1.25 μL CROPDialOut_U6_F (25 μM). The reaction cycled through the following conditions 98°C for 30 sec, (98°C for 15 sec, 69°C for 15 sec, 72°C for 20 sec) x 9 cycles, 72°C for 2 mins. After cleaning the product with 1x SPRI and eluting in 15 μL of H_2_O, it was subsequently used as a template in a second PCR reaction to attach Nextera adapters. 12.5 μL of the SPRI-cleaned PCR product was mixed with 15 μL Phusion High-Fidelity PCR Master Mix, 1.25 μL CROPDialOut_P5_R1 (25 μM), and 1.25 μL of a unique P7 index primer. The reaction was amplified through an additional 9 cycles following the conditions of the first PCR, while adjusting the melting temperature to 57°C. A 0.7x SPRI clean was performed twice to purify the product, eluting in 50 μL H_2_O and 35 μL H_2_O respectively. All primer sequences are listed in **Supplementary Data Table 8**. scRNA-Seq and CITE-seq libraries were sequenced on Illumina HiSeqX to 25,000 reads/cell and 10,000 reads/cell respectively.

### Pre-processing of single-cell data

Expression matrices, representing Unique Molecular Identifier (UMI) counts for both scRNA-seq and CITE-seq data were obtained using the Cumulus (*34*) version 0.14.0 implementation of the CellRanger v.3 workflow with genome reference GRCh38 v3.0.0 and default parameters. Cells with fewer than 200 detected genes or with >18% of detected genes labeled mitochondrial were removed from subsequent analysis. Genes detected in fewer than 200 cells were also removed from further analysis.

### Normalization and integration of single-cell data

scRNA-seq count data was normalized to 1,000,000 total counts per cell (transcripts per million), followed by a natural log transformation. CITE-seq data was normalized according to the formula (*34*):

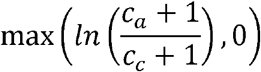

where *c_a_* is the UMI counts for antibody a and *c_c_* is the UMI counts for its corresponding IgG control *c* (**Supplementary Data Table 3**). The two modalities were integrated by concatenation of their normalized count matrices for further analysis.

### Differential expression testing

Differential expression for CITE antibodies across experimental conditions was performed using the Seurat implementation of logistic regression (*35*) with the “ADT” assay and default parameters. Differential expression for transcripts across experimental conditions was performed using MAST (*36*) with default parameters. Differential expression testing relating to Figure 3 was performed on cells with non-targeting sgRNAs only.

### Identification of cellular programs using jackstraw PCA

Jackstraw PCA was implemented according to the algorithm described by Chung and Stoery (*37*). Briefly, expression matrices from each condition were processed separately using MOI = 1 cells only. A subset of 150 features (out of all genes passing QC and all 20 CITE proteins) was randomly sampled, followed by PCA, for 150,000 iterations. This allowed a PC-wise calculation of the synthetic null distribution of PC loadings. PCA was then run on the full expression matrix, to calculate an empirical p value for each feature. A threshold (p < 0.03) was used to assign specific features to each program (PC) to obtain programs of reasonable biological size. The Scikit-learn implementation of PCA was used in all cases (*38*).

### Initial sgRNA assignment to cells

Dial-out PCR sequencing data was processed with custom code to identify cell barcode (CBC) and sgRNA barcode (GBC) pairs. A dictionary was built mapping each CBC-GBC pair to a list of UMIs, each with its associated number of reads. A GBC was assigned to a particular CBC if it exceeded any one of the following thresholds: (**1**) a UMI with more than 200 reads, (**2**) more than 20 total UMIs, or (**3**) a UMI with more than 50 reads that accounts for more than 20% of the total reads for the CBC-GBC pair.

### Computational model for Perturb-CITE-seq analysis

To infer a model of gene regulation from Perturb-CITE-Seq data, following initial sgRNA assignment to cells, we performed feature selection. For each of the three conditions (control, coculture and IFN γ), 1,000 highly variable genes were selected using the Scanpy version 1.4.4 implementation of highly variably gene selection (*39*). In addition, all features from each condition’s 10 most significant Jackstraw PCA programs were included and all 20 CITE antibodies. The union was taken across all conditions resulting in one set of 4,481 features used for each experimental conditions.

Next, for each condition separately, we learned a linear model to predict the effect of each sgRNA on each feature using the Scikit-learn implementation (*38*) of elastic net regularization with the following parameters: l1_ratio = 0.5, alpha = 0.0005, max_iter = 10000.

Fit quality was assessed by correlation of model residuals. Inclusion as covariates of cell quality (defined as the total number of UMIs in each cell) and cell cycle state (assigned with Scanpy’s implementation of scoring cell cycle genes (*39*)) improved the fit. Following the first elastic net regularization, an expectation maximization (EM)-like procedure was run along the columns of the sgRNA assignment matrix to account for false positive sgRNA assignments and sgRNAs that did not perturb their target, using our previously published MIMOSCA framework (*2*). Briefly, each cell is modeled as coming from either a perturbed or unperturbed population. For each sgRNA, the elastic net regularization is rerun with perturbed cells assigned to the unperturbed population. The difference in sum of squares error between the perturbed and unperturbed elastic net predictions was used to estimate to a likelihood that each individual cell was perturbed.

Running this procedure across all sgRNAs effectively transforms the binary sgRNA assignment matrix to a probabilistic estimate of successful perturbation. Following this procedure, elastic net is rerun with the probabilistic sgRNA assignment matrix. The procedure is iterated only once (and thus is not a full EM).

The regulatory matrix fit by this procedure was used to assess concordant effects across sgRNAs with the same target by calculating the pairwise correlation between the regulation profiles for sgRNAs across all features. The Pearson correlation of sgRNAs with different targets was then used to calculate a synthetic null distribution of discordant perturbations, and the pairwise Pearson correlation of sgRNAs with the same target was transformed into an empirical p-value, followed by a Benjamini-Hochberg multiple hypothesis False Discovery Rate (FDR) correction. To aggregate from sgRNAs to the gene level, sgRNAs with correlated activity were mapped to their target gene to construct a sgRNA to target dictionary. A lenient FDR threshold of 0.50 was used due to the large feature space and the sparsity of elastic net regularization. This results in 141, 183, and 181 gene-level targets (of 248 total target genes) for the control, IFNγ, and coculture conditions, respectively. All non-targeting sgRNAs were collapsed to the same target while intergenic sgRNAs were kept separate as individual negative controls.

The sgRNA assignment matrix was mapped to a binary target assignment matrix using the sgRNA to target dictionary for the next iteration of elastic net regularization. An EM-like procedure was then run again according as above, except at the target rather than sgRNA level. A final elastic net regularization was run following the reassignment step.

A permutation test was used to assess the empirical significance of regulatory coefficients as we previously described (*2*). Briefly, for each target, the assignment of cells to targets was randomly permuted, followed by recomputing the linear model. This procedure was run separately for each target, each with 10,000 random permutation, to calculate target-wise null distributions of coefficient values. The coefficients of the model with correct cell assignment are then transformed to empirical p values. For FDR matrices, a Benjamini-Hochberg multiple hypothesis correction is performed target-wise with FDR = 0.25.

### Scoring programs across cells and with perturbations

Programs were scored according to a previously described procedures (*40*)’(*41*). Briefly, expression values are first sorted into 50 expression bins. The expression of a target list of genes is then scored as their average expression less the expression of a control set. The control set is a randomly sampled set of genes with size equal to the number of genes in the target list (unless the target list has fewer than 50 genes, in which case the number of control genes was 50). In the case of programs with both an up- and down-regulated components (*e.g.*, positive and negative loading for the jackstraw programs), the two components are scored separately and the composite score is calculated as the score of the up-regulated component minus the score of the down-regulated component.

The sgRNA to target dictionary (above) was used to assess the effect of perturbations on program scores. Cells were scored according to the procedure above. Cells with sgRNAs mapping to the same target were grouped. Each target had the program score of its cells compared to the score of cells with a non-targeting sgRNA. Welch’s T test was used to test significance for enrichment or depletion of a given program.

### Single gene knock-out in patient-derived cells using nucleofection of Cas9 ribonucleoprotein

Virus free knock-out cell lines of human melanoma cell lines were generated using nucleofection of Cas9 ribonucleoproteins (RNP). Target sequences were derived from the original ICR library used to generate the CROP-Seq library and are listed individually for the perturbed targets in **Supplementary Data Table 8**. To form RNPs, equimolar ratios of crRNA (IDT, Coralville, IA) were incubated with tracrRNA (IDT) at 95°C for 5 minutes in nuclease-free duplex buffer and thereafter cooled to room temperature to from gRNA complexes. Recombinant Cas9 enzyme (MacroLab, UC Berkeley, CA) was mixed with gRNA at 1:10 molar ratio and incubated at 37°C for 15 minutes to form RNP complexes. In the meantime, human melanoma cell lines were detached, washed once with PBS, and 5×10^4^ cells were resuspended in 20 μL SF electroporation buffer prepared with SF supplement (Lonza, Basel, Switzerland). 3 μL RNP complex solution was mixed with the cells and the cells were nucleofected using program DJ-110 on a 4D-Nucleofector (Lonza) with 16-well strips. Cells were immediately recovered in full melanoma media, seeded in 12 well plates and expanded.

### Enrichment of surface protein knockouts using fluorescence activated cell sorting

KO cell lines generated using nucleofection of Cas9 RNP were detached with Accutase (Stem Cell Technologies, Vancouver, Canada) and stained with APC conjugated antibodies targeting B2M (2M2), CD58 (TS2/9), and CD274 (29E.2A3, all Biolegend, San Diego, CA). Unperturbed cells stained with the same antibodies and isotype controls were used as gating controls. Cells negative for B2M, CD58, or CD274 were sorted on an Influx Cell sorter (BD), grown out and banked until further use.

Prior to use in experiments, purity of KO cell lines was assessed using the same clonotypes as above using an Aurora Spectral Analyzer (Cytek Biosciences, Freemont, CA). In addition, loss of MHC class I expression in B2M knock-outs was assessed by staining with FITC-conjugated anti-HLA-A,B,C (W6/32, Biolegend). To maximize surface expression of IFNγ regulated proteins, cells were assessed with and without 16h pretreatment of 10 ng/ml recombinant human IFNγ (Peprotech, Rocky Hill, NJ).

### Competition experiments between labeled parental and KO cells

For mixing studies of single gene knock-outs with wildtype cells, knock-outs were generated in NLS-dsRed target cells and target cells were mixed with unmodified control target cells expressing NLS-BFP. Co-cultures and analyses were performed as described in the miniaturized co-culture assay above. Fold enrichment was calculated as dsRed(viable)/BFP(viable) and normalized to dsRed(viable)/BFP(viable) in conditions without TILs.

### Flow cytometry analysis of surface proteins after IFNγ stimulation

2686 control and CD58 KO cells were stimulated with 1-10 ng/ml IFNγ for 72 hours and then detached with Accutase. Dead cells were labeled using Zombie-NIR (Biolegend) according to the manufacturer’s instructions. Surface antigens were stained on ice using the following antibodies (all Biolegend): HLA-DR-BV421 (L243), HLA-A,B,C-BV605 (W6/32), CD274-BV785 (29E.2A3), and CD58-APC (TS2/9). Cells were fixed in fixation buffer (Biolegend) and analyzed on an Aurora Spectral Analyzer (Cytek Biosciences). The samples were than analysed using FlowJo (FlowJo, Ashland, OR). For quantification of surface proteins. cells were gated based on FSC and SSC, single cells were selected using FSC-A and FSC-H and viable cells were selected using low Zombie-NIR fluorescence. To compare signal intensities of surface markers, geometric mean fluorescence intensity (gMFI) was used and all samples were run in duplicates.

