## Supplementary Material for "Multi-modal pooled Perturb-CITE-Seq screens in patient models define novel mechanisms of cancer immune evasion"

#### Supplementary Figures

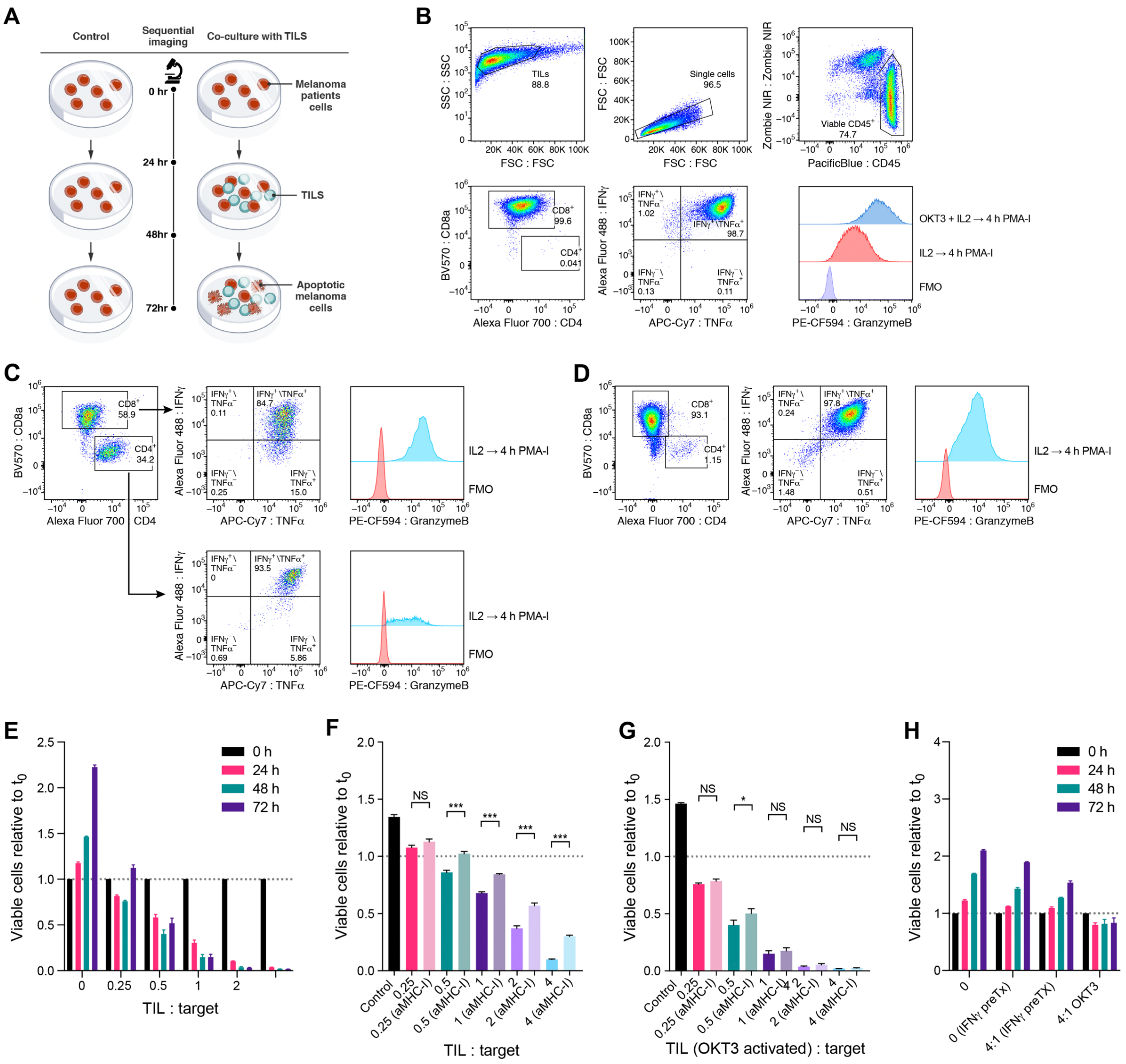


### Figure S1. Establishment of patient derived co-culture model

**(A)** Approach for imaging-based quantification of TIL-mediated killing of melanoma target cells. Plates were imaged at 0, 24, 48 and 72 hours, and viable cell counts were normalized to starting counts to quantify outgrowth of target cells. **(B)-(D)** Sorting and gating strategy to isolate and expand TIL cultures prior to co-culture. **(B)** Flow cytometry plots of TILs grown in recombinant human IL2 or restimulated using immobilized OKT3 for 72 hours and analyzed after 4 hours of

Phorbol-Myristate-Acetate and Ionomycin (PMA-I) treatment. Gating is performed by FSC and SSC, single cells are selected by FSC-A and FSC-H, and viable cells are discriminated by CD45 and Zombi-NIR expression. **(B)** Expanded TILs from tumors 2686 show a pure CD8 population with the ability to induce IFNγ and TNF α, and OKT-3 reactivation leads to an increase in Granzyme-B production compared to TILs grown in IL2 alone. **(C)** Expanded MaMel-134 TILs after 4 hours of PMA-I treatment show $CD4^+$ and $CD8^+$ T cells with the ability to induce IFNγ, TNFα, and Granzyme-B. **(D)** Expanded MaMel-80 TILs after 4 hours of PMA-I treatment are dominated by CD8 T cells with the ability to induce IFNγ, TNFα, and Granzyme-B. **(E)-(H)** Impact of time, dose, IFNγ pre-treatment, MHC-I blocking antibodies, and OKT3 on TIL mediated killing in the co-culture system from patient 2686. Ratio of viable cancer cells (y axis, relative to t0) in co-cultures: (**E**) after different time points of co-culture at increasing TIL:cancer cell ratios (x axis), where TILs were restimulated with immobilized OKT3 for 72h prior to co-culture; (**F**) after 48h of co-culture, where cancer cells were pre-treated with 1 ng/ml IFNγ for 16 hours, TILs were not restimulated prior to co-culture, and cultures were with or without MHC-I blocking antibody; (**G**) after 48 hours of co-culture as in **(F)** but using OKT3-reactivated TILs. Experiments were performed in triplicates and is representative of two consecutive experiments. ns: $p>0.05$; * $p<0.05$; ** $p<0.005$; *** $p<0.001$ Two-way ANOVA with Tukey *post hoc* test. Error bars: Mean ±SD . (**H**) Specificity of IFNγ pre-treatment approach. Ratio of viable allogenic cancer cells (y axis, relative to t0) in different culture conditions with or without IFNγ pre-treatment (x axis) grown from 0 to 72 hours (color bars) with 2686 TILs with or without prior reactivation with OKT3. Error bars: Mean ±SD

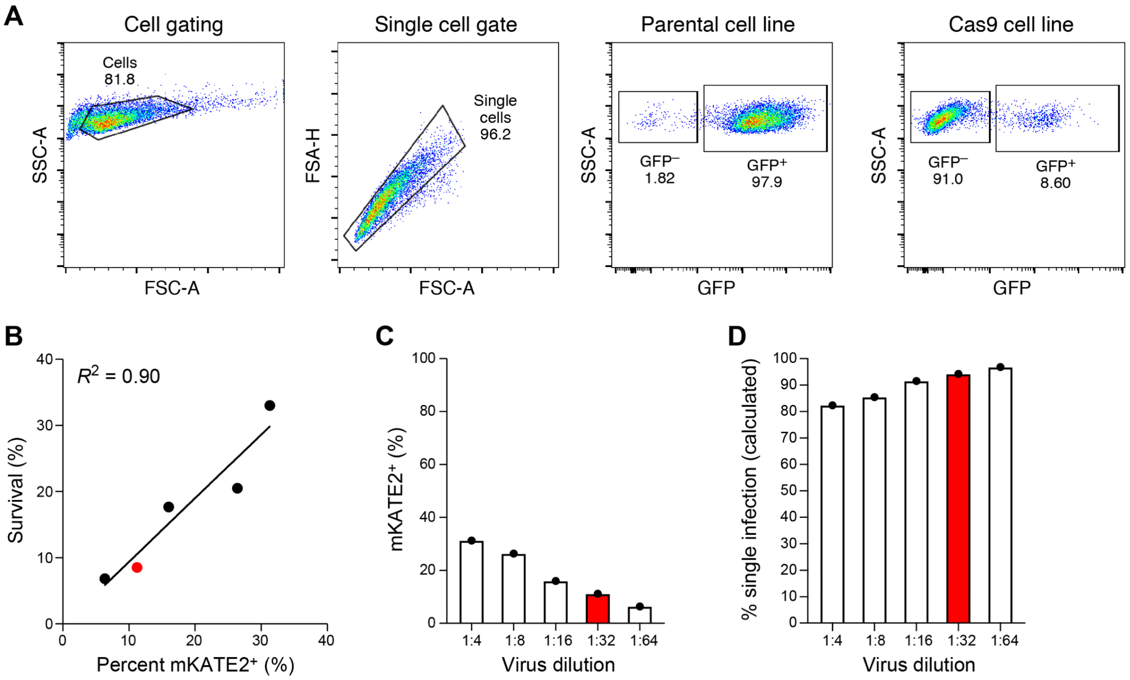


**Figure S2. Generation of Cas9 transgenic patient derived lines and sgDNA library titration**

**(A)** High Cas9 activity in Cas9 transgenic line. Flow cytometry of EGFP levels Cas9 transgenic and parental melanoma cells from patient 2686 transduced with lentivirus encoding EGFP and an EGFP targeting sgRNA at MOI<1 and selected using puromycin. **(B)-(D)** Transduction of sgDNA lentiviral library to Cas9 transgenic line. **(B)** Proportion of mKATE2$^+$ cells prior to selection (*x axis*) and survival after puromycin selection (*y axis*) in 2686 melanoma Cas9 transgenic cells transduced with the ICR library. Line: Linear regression, Pearson $R^2$=0.90. **(C)** Percentage of mKate2$^+$ cells (y axis) in 2686 melanoma Cas9 transgenic cells transduced with the ICR library at virus dilutions (x axis). Red: Dilution used for the Perturb-CITE-seq screen. **(D)** Proportion of cells estimated to be infected by one virus (y axis) at different dilutions of the ICR library (x axis). Red: Dilution used for the Perturb-CITE-seq screen.

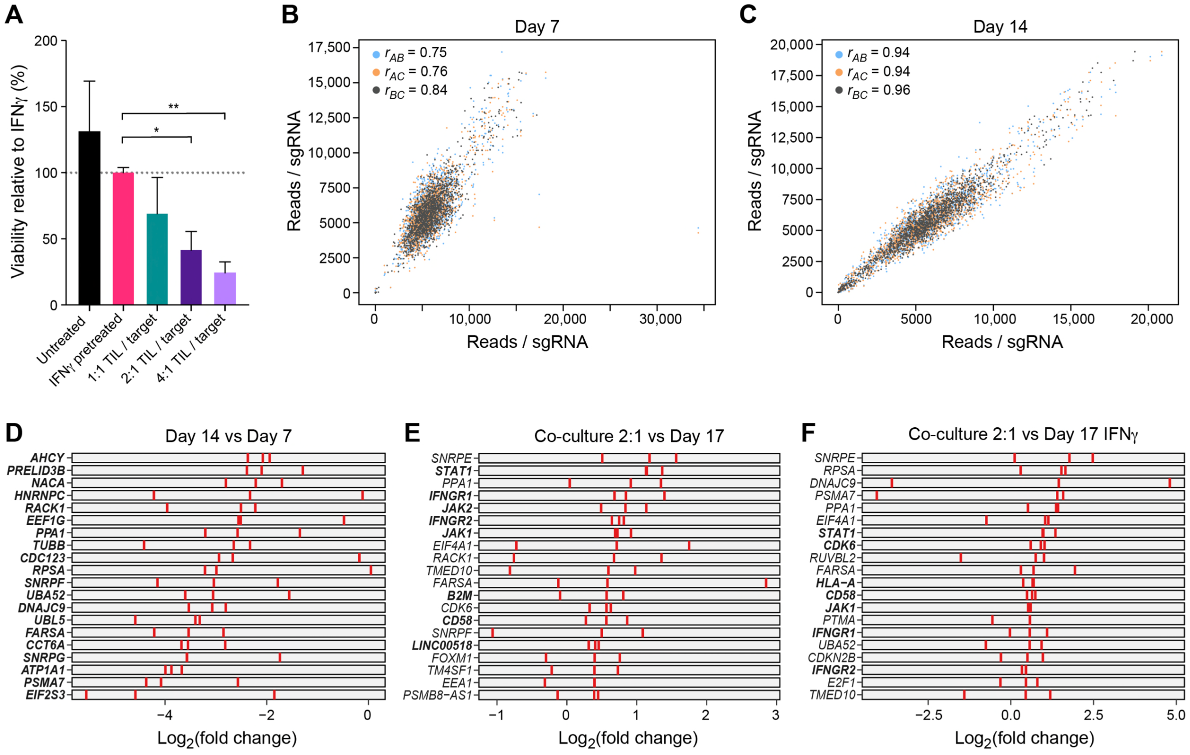

**Figure S3**. **CRISPR-Cas9 viability screen in the co-culture system**

**(A)** Dose-response killing in the co-culture experiment validates target killing range. Percent of surviving cells relative to IFNγ pretreated target cells (y axis) in different co-culture conditions (x axis) from a plate run in parallel to the virability and Perturb-CITE screens, with triplicate wells for each condition. ns: p>0.05; * p<0.05; ** p<0.005; *** p<0.001, one-way ANOVA with Dunnet *post-hoc* test. **(B)(C)** Screen reproducibility across triplicates. Number of reads detected (*x*, *y* axis) for each sgDNA (dots) when comparing each pair within triplicate experiments (color legend) in pre-treated day 7 **(B)** or day 14 **(C)**. Pearson correlation coefficients are noted in the color legend. **(D)-(F)** Identification of essential genes and genes affecting resistance to TIL mediated killing. Relative depletion (log2(FC), x axis) for each individual sgDNA (red bar) of the top20 target genes by MAGeCK analysis (rows) (n=3 sgDNAs/target gene, **Methods**) on day 7 (without TILs, to recover essential genes, (**D**)), day 17 of 2:1 TIL:cancer cells co-cultures, comparing control cells (**E**), or day 17 of IFN*γ* treated cells ((**F**), no co-culture). Bold lettering indicates significantly enriched / depleted target.

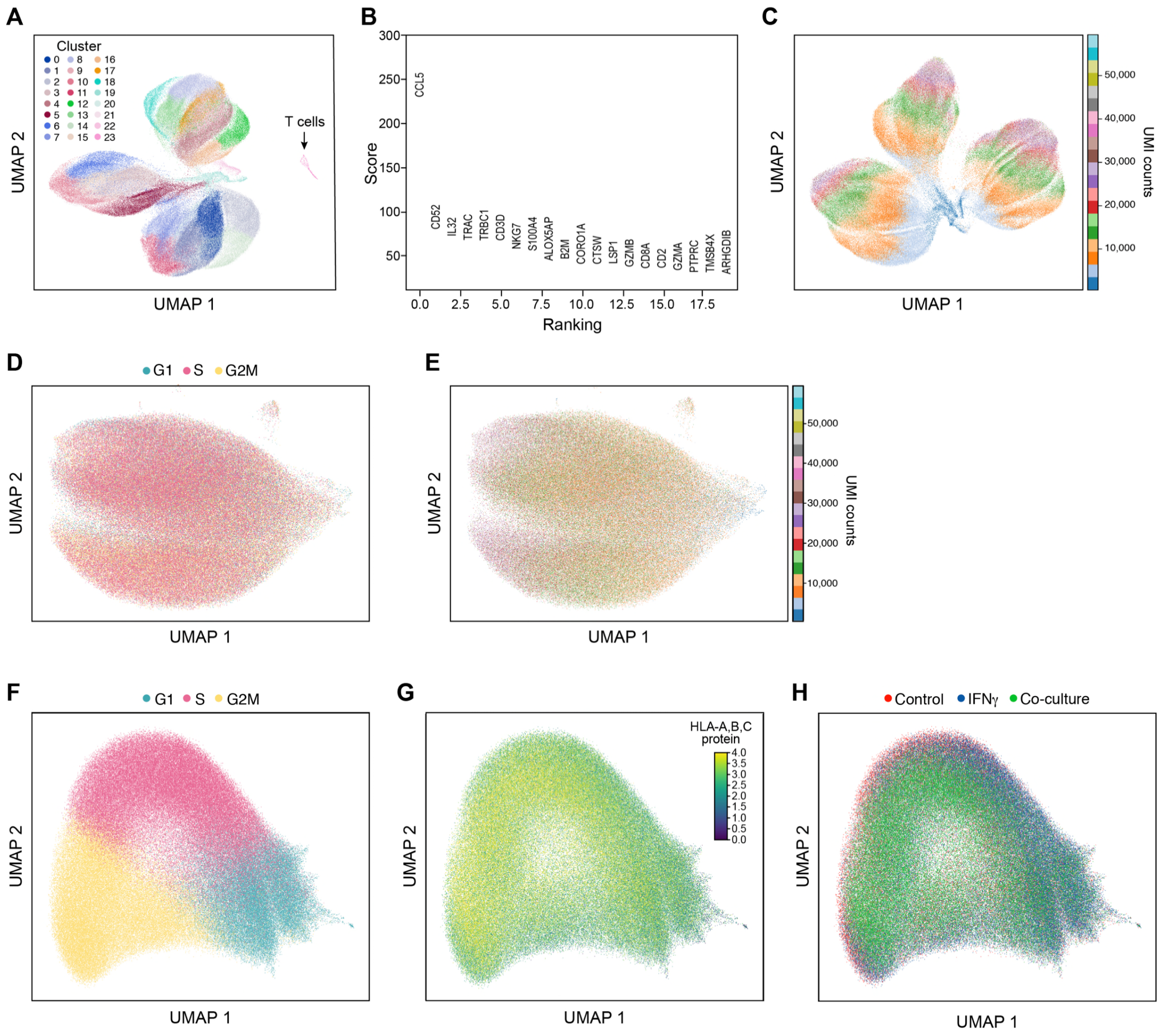


**Figure S4. Characterization of different immune pressures by single-cell RNA and protein profiles**

**(A)(B)** Removal of profiled T cells. **(A)** UMAP embedding of single cell RNA-Seq profiles from the Perturb-CITE-Seq screen, colored by unsupervised cluster assignment (*42*) (**Methods**). **(B)** A permutation test was used to score marker genes associated with each cluster shown in (A) (*39*). Score (y axis, permutation test, **Methods**) of marker genes (x axis) associated with the distinct cluster marked by an arrow in (**A**), include canonical T cell markers. **(C)** Cell complexity affects scRNA-Seq profiles. UMAP embedding of scRNA-Seq profiles of cancer cells only, colored by

UMI count bins. **(D)(E)** CITE profiles of 20 cell surface proteins do not reflect cell cycle phases **(D)** or UMI count **(E)**. UMAP embedding of cells (dots) by CITE-seq profiles (dots) colored by cell cycle phase, as scored from scRNA-Seq of the cells, and **(E)** UMAP of cells by count bins (indicated in legend). **(F)-(H)**. Limited relation between the cell cycle and immune pressure or phenotype. UMAP embedding of cells (dots) based only on RNA expression on cell cycle genes colored by (**F**) cell cycle phase based on the cell's RNA profile; (**G**) HLA protein levels from the CITE signal of the cell; or (**H**) condition.

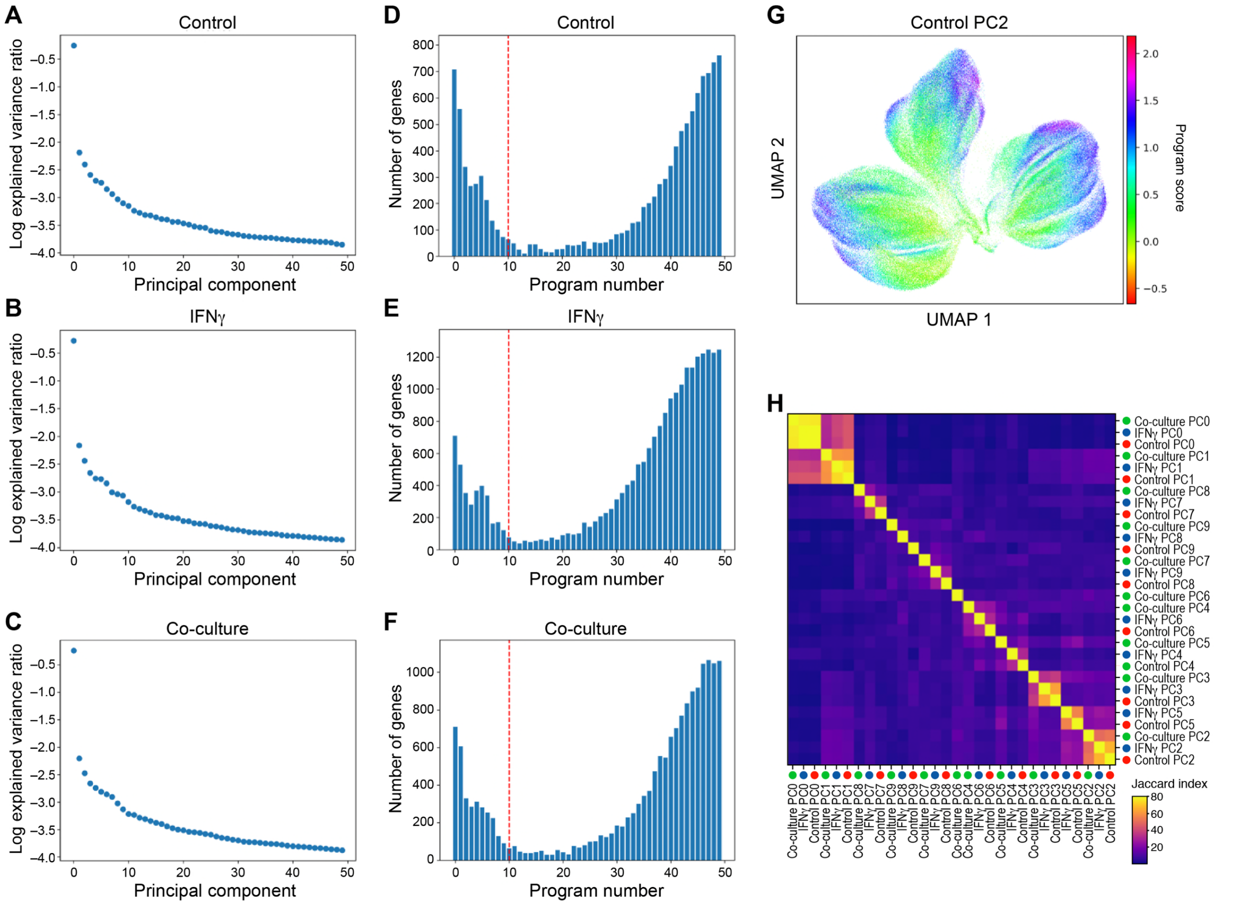


**Figure S5. Learning expression programs in different conditions**

**(A)-(F)** Identification of programs by jackstraw PCA in each condition. **(A)-(C)** Explained variance (y axis) by each principal component (x axis) for PCA performed on control **(A)**, IFN$\gamma$-treated **(B)**, or co-culture **(C)** Perturb-CITE-Seq data. **(D)-(F)** Number of features (y axis) for each jackstraw program (x axis) for models learned on control **(A)**, IFN$\gamma$-treated **(B)**, or co-culture **(C)** Perturb-CITE-Seq data. Dotted red line: cutoff for programs considered in further analysis. **(G)** G2M program learned from control dataset. UMAP embedding of cells (dots) by scRNA-seq profiles, with cells colored by the gene set score (color bar) (**Methods**) of a G2M cell cycle control program (compare to **Figure 3F**) (**Methods**). **(H)** Identifying related programs across conditions. Jaccard index (color bar) for each pair of programs across all 30 programs (rows).

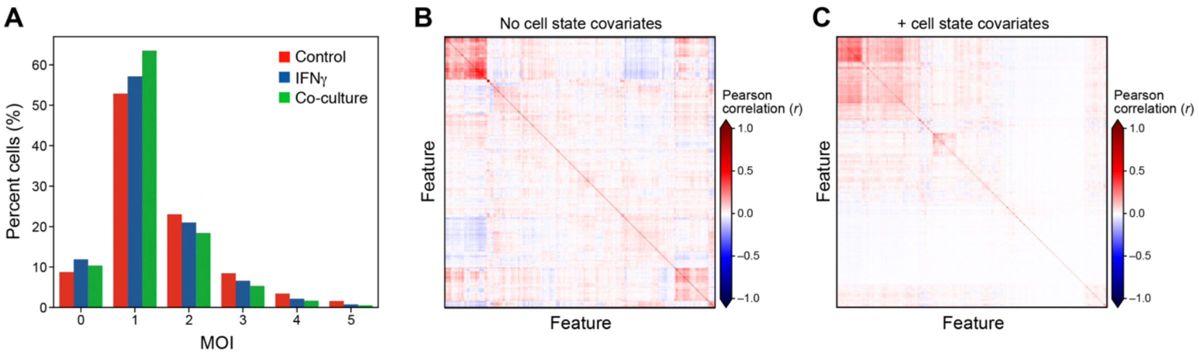


**Figure S6. Addressing cell cycle and complexity covariates by the Perturb-CITE-Seq model and impact of targeting *vs*. non-targeting guides**

**(A)** Estimated Multiplicity of infection (MOI). Distribution of cells (%, y axis) at different estimated MOI (x axis) in each experimental condition (color legend) as determined from the guide dictionary (**Methods**). **(B)(C)** Improved model fit following accounting for cell state as a covariate. Pearson correlation (color bar) between the residuals from the linear model fit for each regulated feature (columns) from models learned without (B) or with (C) cell state covariates accounting for the cell cycle and cell complexity.

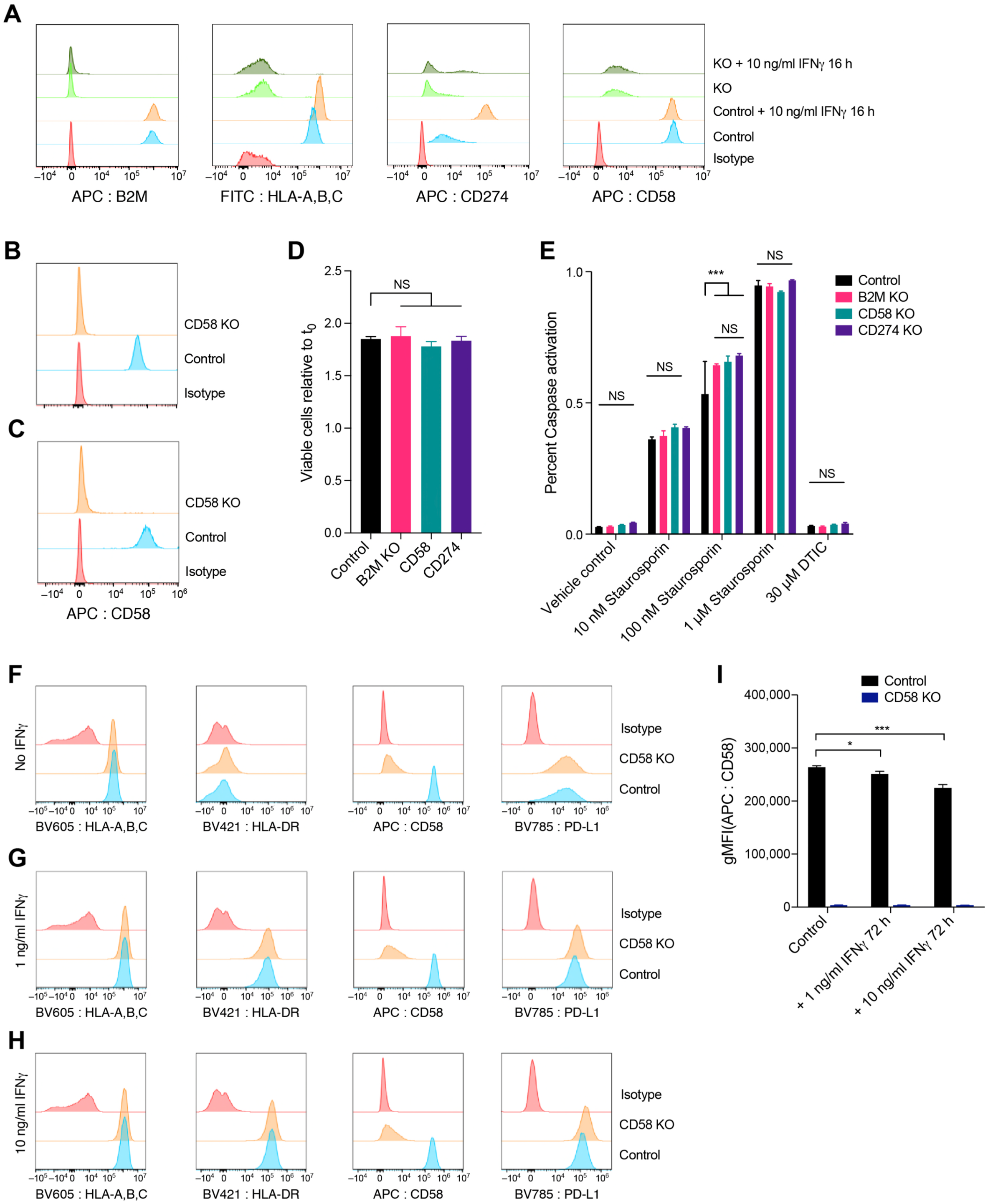


**Figure S7. Role of CD58 in resistance to T cell mediated killing and regulation of PD-L1**

**(A)** Validation of CRISPR-Cas9 KO in patient derived melanoma lines. Distribution of fluorescent intensity by flow cytometry of anti-CD58 antibody (APC-CD58, **(A)-(C)**), as well as B2M (APC-B2M, **(A)**), MHC-I (FITC-HLA-A,B,C, **(A)**), CD274 (APC-CD274, **(A)**), and corresponding isotype controlled stained respectively in control (unperturbed) cells and in CD58KO, B2MKO, and CD274 KO melanoma cells from patient 2686, with or without pre-stimulation with IFN γ for 16 hours (**A**) and in control and CD58KO cells from MaMel-134 (**B**) and MaMel-80 (**C**) melanoma cells. **(D)** Comparable growth of control and KO cells. Ratio of viable cells relative to timepoint 0 (y axis) for control and B2M KO, CD58 KO or CD274 KO melanoma cells from patient 2686 (x axis). ns: $p>0.05$; * $p<0.05$; ** $p<0.005$; *** $p<0.001$; one-way ANOVA with Tukey *post hoc* test. Error bars: Mean ±SD. **(E)** Comparable induction of apoptosis in response to Staurosporin and resistance to DTIC in control and KO melanoma cells. Percent of cells inducing Caspase 3/7 (*y* axis) in control and B2M KO, CD58 KO or CD274 KO melanoma cells (color code) from patient 2686 in different treatment conditions (x axis). ns: $p>0.05$; * $p<0.05$; ** $p<0.005$; *** $p<0.001$; two-way ANOVA with Tukey *post hoc* test. Error bars: Mean ±SD. **(F)-(H)** CD58 perturbation in co-culture does not affect B2M and HLA expression at the RNA and protein level but induces CD274. Distribution of fluorescent intensity by flow cytometry (corresponding to **Figure 5H-5J**) of MHC Class I and II, CD58, and CD274 (PD-L1) in parental (control) and CD58 KO lines at baseline (**f**) and after 72 hours of stimulation with either 1 ng IFNγ (**G**) or 10 ng IFNγ (**H**).

**Supplementary Data Table legends**

**Supplementary Data Table 1.** Targeting and control sgRNA sequences and their respective gene target.

**Supplementary Data Table 2.** CRISPR-Cas9 viability screen.

**Supplementary Data Table 3.** CITE-Seq antibodies used in this study.

**Supplementary Data Table 4.** Differential expression of RNA and protein

**Supplementary Data Table 5.** Jackstraw programs.

**Supplementary Data Table 6.** Co-functional modules and co-regulated programs from Perturb-CITE-Seq.

**Supplementary Data Table 7.** CROP-seq vector cloning.

**Supplementary Data Table 8.** Primers used in this study.
